# EXTENDED TUDOR-DOMAINS of the piRNA BIOGENESIS PATHWAY HAVE RNA-SPECIFIC NUCLEASE ACTIVITY

**DOI:** 10.1101/2021.10.25.465661

**Authors:** Neha Dhimole, Susanne zur Lage, Wilfried Klug, Teresa Carlomagno

## Abstract

piRNAs are essential for transposon repression and protecting the germline from deleterious mutations. piRNA biogenesis comprises a primary and secondary pathway, and involves PIWI clade argonaute proteins and ancillary factors. Secondary piRNA biogenesis is tightly coupled to transposon repression. It requires processing of the 3’ end of pre-piRNA during an amplification loop by an as yet unidentified endonuclease. Here, using crystallography, and biochemical assays, we discover that the *Drosophila* Qin protein, which is a critical member of the core amplification complex, has endonuclease activity. Qin contains five extended Tudor domains, which had been proposed to recognize methylated ligands. Instead, we show that these domains act as RNA-specific nucleases. This supports a role for Qin in the 3’ end processing of Ago3-bound pre-piRNAs. Extended Tudor domains are frequent in piRNA-processing proteins, suggesting that the uncovered nuclease activity of this protein fold may be key to understanding the piRNA biogenesis.

## INTRODUCTION

Small regulatory RNAs are widely present in both plants and animals, where they act at the transcriptional or post-transcriptional level to regulate important cellular processes. Regulatory RNAs include microRNAs, small interfering RNAs (siRNAs) and Piwi-interacting RNAs (piRNAs). They all associate with argonaute-family proteins and direct them for specific nuclease action. Specifically, piRNAs interact with PIWI clade argonaute proteins to suppress transposons in the animal germline.

Transposons are parasitic genetic elements that can move within or between genomes: in this process, they may disrupt gene integrity and introduce mutations (Levin & Moran, 2011). Over geological time scales, transposons have been major drivers of evolution, as testified by their high abundance in mammalian genomes, which is 33-50% (Lee, Ayarpadikannan, & Kim, 2015; Levin & Moran, 2011). On the other hand, transposons represent a threat to an individual genome’s integrity and need to be actively suppressed. In germline cells, failure to suppress transposons can lead to defects in gametogenesis and infertility in offspring (Girard, Sachidanandam, Hannon, & Carmell, 2006; Ozata, Gainetdinov, Zoch, O’Carroll, & Zamore, 2019). In animal gonads, transposon suppression depends on non-coding RNAs of 24-32 nucleotides in length, called the PIWI-interacting RNAs (piRNAs), which guide the PIWI clade argonaute proteins to cleave transposon transcripts (Aravin et al., 2006; Aravin, Hannon, & Brennecke, 2007; Czech & Hannon, 2016; Girard et al., 2006; Lau et al., 2006). As an indispensable tool to maintain genomic integrity, piRNAs are conserved in the entire animal kingdom (Grimson et al., 2008; Ozata et al., 2019).

The biogenesis of piRNAs differs from that of all other non-coding RNAs. It begins at specific genomic loci, the piRNA clusters, whose transcription yields precursor piRNAs (pre-piRNAs) (Brennecke et al., 2007; Siomi, Sato, Pezic, & Aravin, 2011). The long single-stranded pre-piRNA sequences are transported to the nuage, a specialized cytosolic non-membranous perinuclear structure, where they are further processed via primary and secondary processing pathways to yield functional piRNAs, that repress transposons. Despite much efforts in studying both the primary and secondary processing pathways, many steps of piRNA biogenesis are still poorly understood.

In the secondary pathway, piRNA production is tightly coupled to transposon silencing through a transposon-sequence-based piRNA amplification loop, commonly called the *ping-pong cycle* (Brennecke et al., 2007; Czech & Hannon, 2016). Two essential members of the cycle are the PIWI clade argonaute proteins Ago3 and Aubergine (Aub). Ago3 binds to sense piRNA sequences, recognizes the complementary sequence of a pre-piRNA transcript and cleaves it to generate antisense piRNA (Brennecke et al., 2007). This is then transferred to Aub. The Aub– mature-piRNA complex recognizes and cleaves the sense transposon transcript, thereby generating a new copy of sense piRNA and reinitiating the cycle (Brennecke et al., 2007; Czech & Hannon, 2016; Czech et al., 2018; Siomi et al., 2011). Slicing by Ago3 and Aub generates the 5’ end of the antisense and sense piRNA. The 3’ end trimming and piRNA maturation, which are essential to obtain functional piRNAs, are more complex processes that require the action of one or more yet-unknown nucleases. Reasonably, the still undiscovered nuclease function may be harbored by one of the many additional factors involved in secondary piRNA biogenesis (Czech & Hannon, 2016).

Qin (also called kumo) is an essential protein of the ping-pong cycle. It comprise an N-terminal RING domain followed by five Tudor-like domains (Anand & Kai, 2012; Z. Zhang et al., 2011) (Fig. 1A). Qin knockouts result in developmental disorders such as fused or missing dorsal appendages in *Drosophila* embryos and failure to develop into fully functioning adults. On the molecular level, Qin deletion results in a futile homotypic (Aub-Aub) ping-pong cycle – as opposed to the physiological heterotypic (Aub-Ago3) cycle – leading to increased sense piRNA levels and a concomitant decrease in antisense piRNAs. This homotypic cycle fails to repress transposons, which in turn leads to DNA damage (Zhang et al., 2011). The mammalian Qin homologue TDRD4/RNF17 (ring finger protein 17) is essential in the mammalian germline and its deficiency results in infertility in mice (Pan et al., 2005).

**Figure 1:**
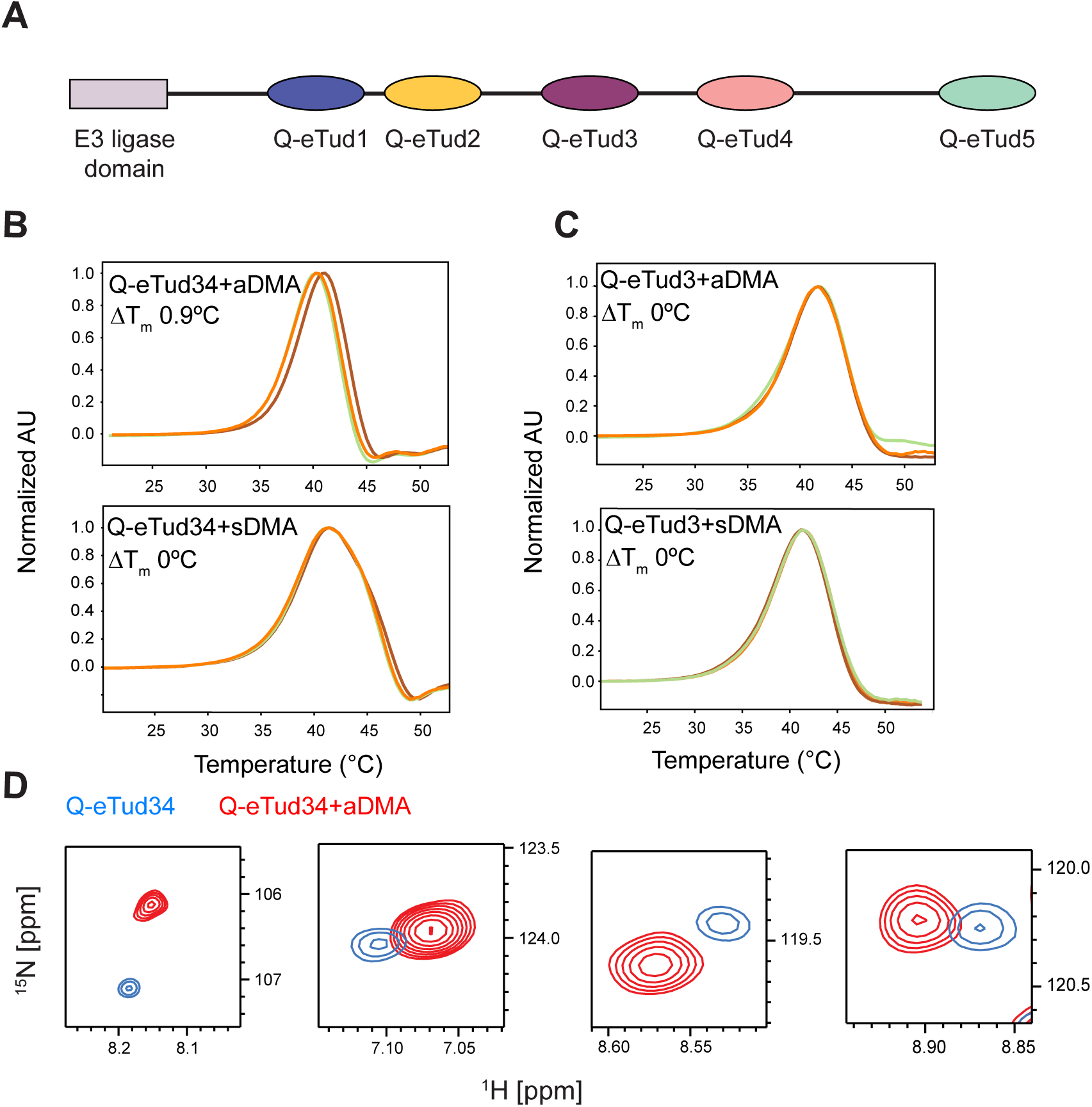
Q-eTud4 interacts with aDMA. **A**. Domain composition of Qin. **B, C**. Thermal Shift Assays (TSA) of Q-eTud34 (B) and Q-eTud3 (C) showing the protein melting temperature (T_m_) in the presence of increasing concentrations of either aDMA or sDMA (green, orange and red curve correspond to the addition of 0, 1 and 10 mM ligand, respectively. **D**. Expansion of four peaks of the NMR ^1^H-^15^N HSQC spectrum of 50 μM Q-eTud34 in the presence (red) and in the absence (blue) of 2.5 mM aDMA. The spectra were acquired at 293 K and 850 MHz ^1^H frequency. The full spectra are shown in Fig. S1.

Tudor domains are present in many of the proteins participating in the ping-pong cycle (Ku & Lin, 2014; Sato, Iwasaki, Siomi, & Siomi, 2015). These domains are known to recognize methylated arginine or lysine (Chen, Nott, Jin, & Pawson, 2011); in agreement, several Tudor domains of proteins active in piRNA-biogenesis, such as *Drosophila* Papi, *Drosophila* eTUD and murine TDRD1, have been shown to recognize sDMA marks in disordered regions of PIWI, Aub and Mili, respectively (H. Liu et al., 2010; Mathioudakis et al., 2012; Y. Zhang et al., 2018). As a consequence, some of the Tudor domain containing proteins have been proposed to act as a molecular scaffolds that may recruit RNA-processing factors to a common platform (H. Y. Huang et al., 2011; L. Liu, Qi, Wang, & Lin, 2011; Mathioudakis et al., 2012). Because both Ago3 and Aub carry symmetrically dimethylated arginine residues (sDMA) in their terminal unstructured regions (Kirino et al., 2009; Vagin et al., 2009), and Qin is required for association of Ago3 with Aub in *Drosophila* (Anand & Kai, 2012; Z. Zhang et al., 2014; Z. Zhang et al., 2011), it is tempting to speculate that Qin functions as a molecular scaffold too.

This hypothesis has been challenged by Xiol et al., who isolated a transient piRNA amplification complex from *Bombyx mori* (*B. mori*) ovarian cell lines, containing antisense piRNA, sense transposon transcripts, Ago3, Siwi, (the *B. mori* Aub homolog), Vasa (a DEAD-box RNA helicase that uses ATP hydrolysis to transfer the newly formed 5’ end of the piRNA from Aub to Ago3) and Qin, and showed that the formation of this complex does not depend on the presence of sDMA markers in Ago3 and Aub (Xiol et al., 2014). In another study, Qin and Siwi were found to co-purify with Snp-E, and this interaction was demonstrated to depend on RNA binding (Andress et al., 2016; K. M. Nishida et al., 2018). In summary, despite the clear phenotype of Qin knockouts and strong evidence of its occurrence in piRNA-biogenesis *in vivo*, the role of Qin in the ping-pong cycle has remained ambiguous.

Here we show that four of the five extended Tudor domains (Q-eTud) of Qin do not bind to post-translational protein modifications. Only Q-eTud4 recognizes asymmetrically dimethylated arginine (aDMA), but not any aDMA-containing Vasa peptide. Instead, Q-eTud1–4 have endonuclease activity, with a strong preference for cleaving single stranded RNA. We determine the atomic structure of Q-eTud3, which adopts an extended Tudor fold, with the Tudor domain intercalated in the SN domain. By structural-based mutagenesis, we demonstrate that the endonuclease activity is harbored in the SN domain of the eTuds. Our findings, together with the Qin knockout phenotype described in the literature, support a function for Qin in the ping-pong cycle as the RNase that processes the 3’ end of piRNA precursors. The frequent occurrence of eTuds in non-coding RNA processing pathways, such as piRNA and micro-RNA (miRNA), suggests a more general role of this protein fold in RNA metabolism.

## RESULTS

### Only Qin eTud4 recognizes methylated ligands

To determine whether the Tudor domains of Qin bind methylated arginine side chains, we expressed and purified the following constructs: Qin^959-1195^ (Qin eTud3, Q-eTud3); Qin^1648-1857^ (Qin eTud5, Q-eTud5), Qin^515-934^ (Qin eTud1–2, Q-eTud12) and Qin^959-1515^ (Qin eTud3–4, Q-eTud34) (Fig. 1A). Q-eTud1, Q-eTud2 and Q-eTud4 were not soluble as isolated domains. A thermal shift assay (TSA) was established to assess the binding of purified Qin constructs to various methylated ligands, such as sDMA, aDMA, mono-methylated lysine (MML), di-methylated lysine (DML) and tri-methylated lysine (TML); in TSA a binding event is detected as a change in the melting temperature (Tm) of the protein.

Qin’s eTuds did not show a significant Tm change in the presence of any of the ligands, with the exception of Q-eTud34, whose Tm increased by 0.9 °C upon addition of 10 mM aDMA (Fig. 1B and Table S1). Since Q-eTud3 did not bind to aDMA, we hypothesized that Q-eTud4 was responsible for the recognition of the ligand in the Q-eTud34 construct (Fig. 1C and Fig. S1B). This hypothesis was confirmed by NMR spectroscopy, where ^1^H,^15^N HSQC spectra of Q-eTud3 and Q-eTud34 were used to detect aDMA binding by monitoring chemical shift perturbations (CSPs) upon addition of excess concentrations of ligand: CSPs were observed in the presence of aDMA for the peaks of Q-eTud4 in Q-eTud34 but not for the peaks of Q-eTud3 (Fig. 1D and Fig. S1).

Among the proteins involved in the ping-pong cycle, only Vasa carries aDMA marks in its N-terminal region (Kirino et al., 2010). Thus, we designed ten 13-amino-acid-long peptides derived from Vasa and carrying the aDMA modification at the central position (Table S2). Screening with TSA failed to reveal any binding of Q-eTud34 to any of the Vasa-derived peptides, probably because we could not reach a peptide concentration higher than 1.2 mM. Similarly, Q-eTud34 did not bind to the sDMA-containing peptides derived from Aub, Vasa and Piwi (Table S2). Finally, we tested a peptide sequence derived from Q-eTud3 containing a GRG motif as putative dimethylation site (Table S2). The interaction of this peptide with Q-eTud4 could function to place the Q-eTud3 and Q-Tud4 domains in a specific orientation to each other. However, also this peptide was unable to bind Q-eTud34.

Also Q-eTud12, Q-eTud3 and Q-eTud5 failed to recognize sDMA and aDMA ligands in the context of peptides derived from Aub, Ago3, Piwi, Vasa and Qin, as well as signatures of mRNA transcribed by RNA-polymerase II, such as 7-methyl-guanosine triphosphate (5’-cap), and cellular RNA modifications, such as 5-methyl-cytosine (5mC) and 6-methyl-adenosine (6mA). Exceptions to this were Q-eTud3, which recognized 7mG in isolation but not in the Q-eTud34 construct, and Q-eTud34 in combination with 5mC (Table S1).

In summary, Q-eTud1–3 and Q-eTud5 do not bind to aDMA, sDMA or any aDMA/sDMA-modified peptides. Q-eTud4 recognizes high concentrations of aDMA but fails to bind to 10 times smaller concentrations of Vasa-derived peptides containing the aDMA mark. All in all, these data suggest that Q-eTud1–5 have a non-canonical function that differs from the recognition of post-translational modifications.

### Qin’s five eTuds belong to the extended Tudor family

To generate hypotheses for the function of the Q-eTuds, we solved the crystal structure of Q-eTud3 (Table 1) and probed the structure of Q-eTud5 by means of small-angle X-ray scattering (SAXS). During the purification of Q-eTud3 via size-exclusion chromatography we noticed that a portion of the protein formed multimers (Fig. S2); multi-angle light scattering experiments (MALS) identified them as dimers, with a molecular weight of ∼60 kDa. For crystallization, we selected the peak corresponding to the monomeric form of the protein.

**Table 1.**
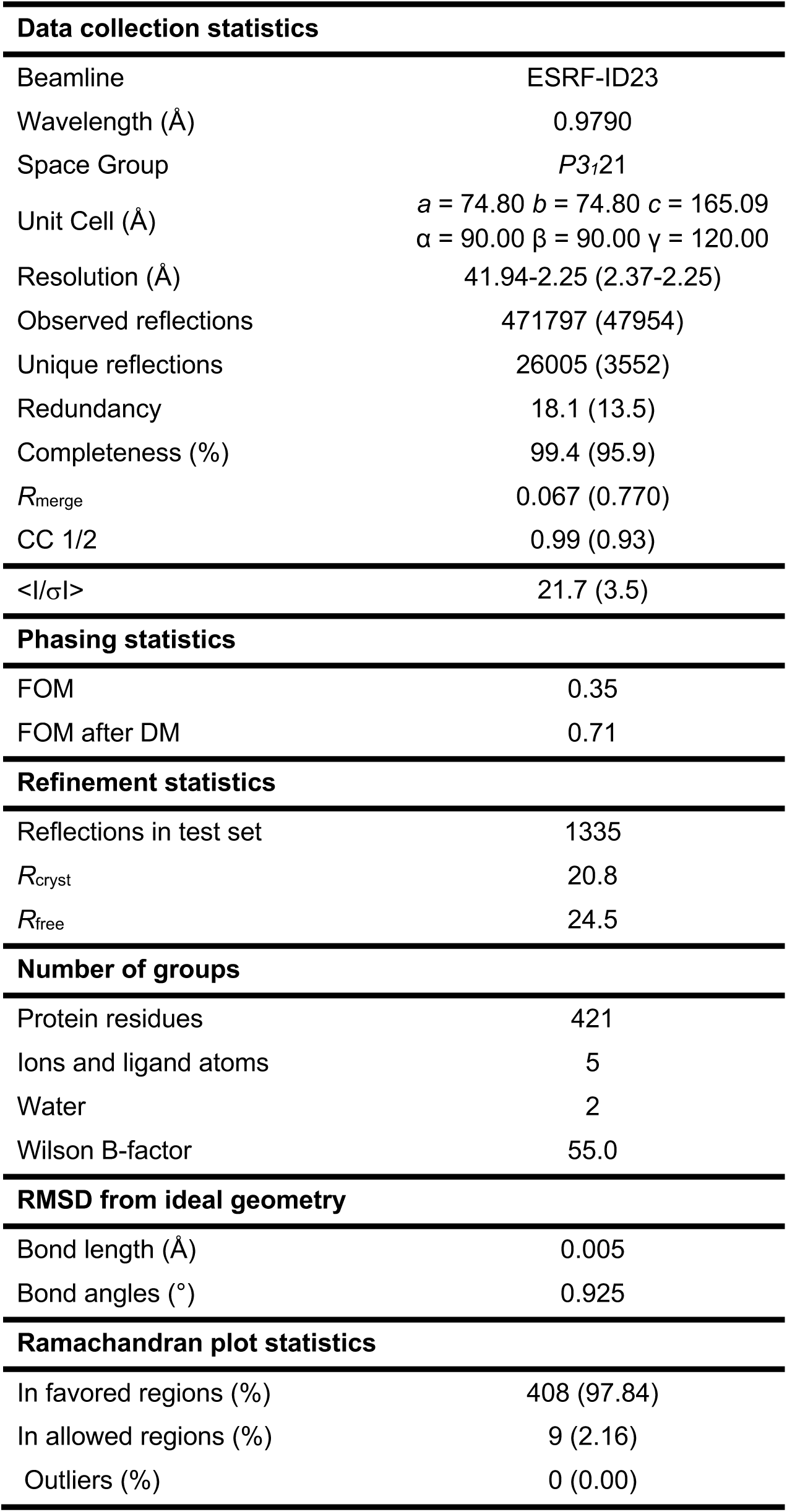
Crystallization data and refinement statistics for Q-eTud3. Values in parentheses are for the highest resolution shell.

| **Data collection statistics** |  |
| --- | --- |
| Beamline | ESRF-ID23 |
| Wavelength (Å) | 0.9790 |
| Space Group | *P*3<sub>1</sub>21 |
| Unit Cell (Å) | *a* = 74.80 *b* = 74.80 *c* = 165.09<br>$\alpha$ = 90.00 $\beta$ = 90.00 $\gamma$ = 120.00 |
| Resolution (Å) | 41.94-2.25 (2.37-2.25) |
| Observed reflections | 471797 (47954) |
| Unique reflections | 26005 (3552) |
| Redundancy | 18.1 (13.5) |
| Completeness (%) | 99.4 (95.9) |
| *R*<sub>merge</sub> | 0.067 (0.770) |
| CC 1/2 | 0.99 (0.93) |
| <\|*I*\|/$\sigma$\|> | 21.7 (3.5) |
| **Phasing statistics** |  |
| FOM | 0.35 |
| FOM after DM | 0.71 |
| **Refinement statistics** |  |
| Reflections in test set | 1335 |
| *R*<sub>cryst</sub> | 20.8 |
| *R*<sub>free</sub> | 24.5 |
| **Number of groups** |  |
| Protein residues | 421 |
| Ions and ligand atoms | 5 |
| Water | 2 |
| Wilson B-factor | 55.0 |
| **RMSD from ideal geometry** |  |
| Bond length (Å) | 0.005 |
| Bond angles (°) | 0.925 |
| **Ramachandran plot statistics** |  |
| In favored regions (%) | 408 (97.84) |
| In allowed regions (%) | 9 (2.16) |
| Outliers (%) | 0 (0.00) |

Q-eTud3 adopts an extended Tudor fold comprising of an SN domain intercalated by a Tudor domain (Fig. 2). eTud domains have been found in many other proteins involved in nucleic acids metabolic pathways, including TUD and TDRD in piRNA biogenesis, and Tudor-SN/P100 in miRNA decay (Elbarbary et al., 2017; H. Y. Huang et al., 2011; C. L. Li, Yang, Chen, & Yuan, 2008; Mathioudakis et al., 2012; Kazumichi M. Nishida et al., 2009; K. M. Nishida et al., 2018; Ren et al., 2014). The SN fold originates from the Staphylococcal Nuclease protein, which falls under the superfamily of OB domains. The SN domain of Q-eTud3 is formed by 6 β-strands and 2 α-helices and differs from the SN domains of P100 in that it shows an extended β-strand–loop– β-strand motif extending from β-strand 9 (the numbering is as in Q-eTud3, Fig. 2A). In addition, the C-terminal helix present in the SN domain of both P100 and the Staphylococcal Nuclease was absent in the construct of Q-eTud3 used for crystallization (Fig. 2B)

**Figure 2.**
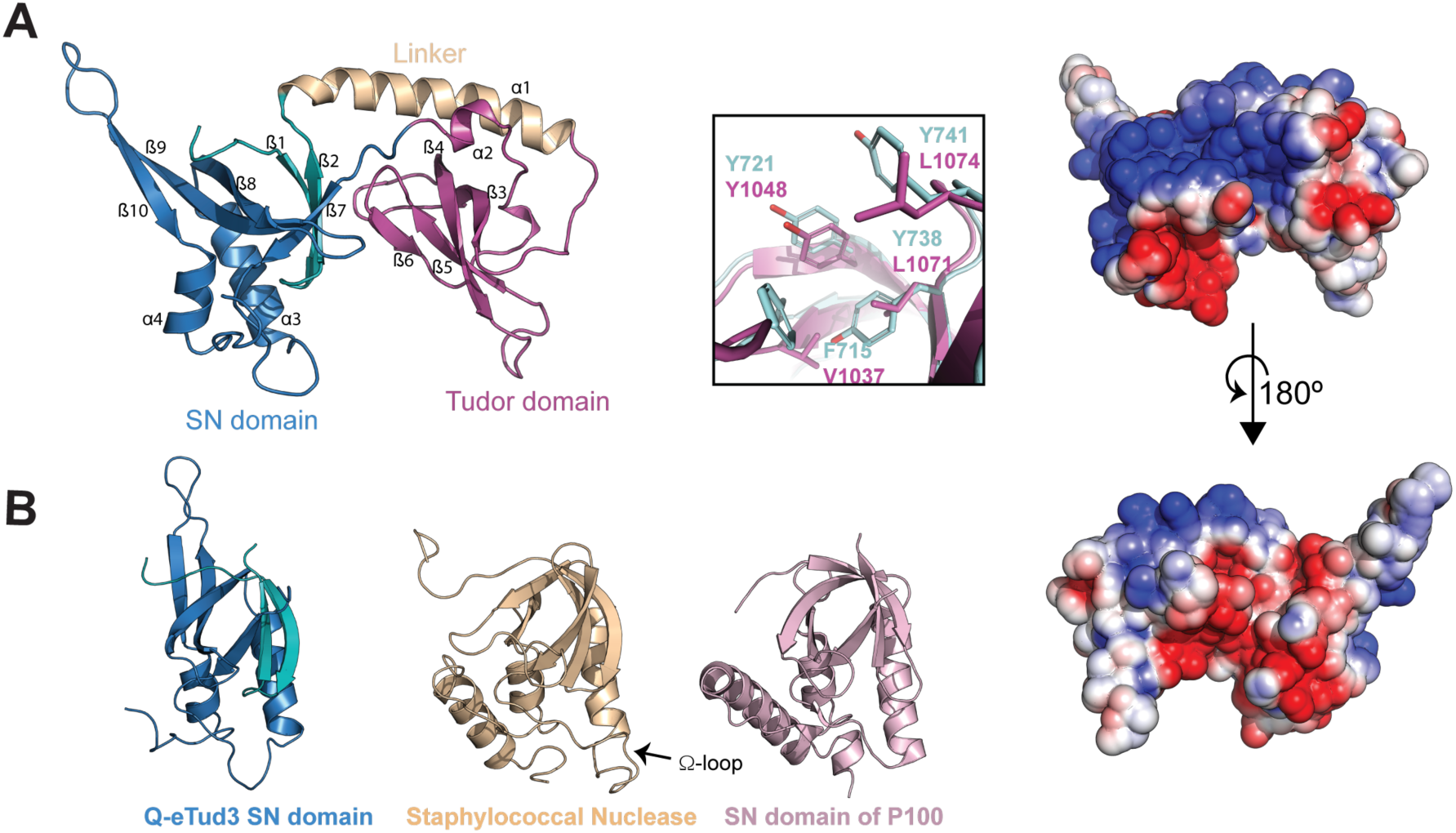
X-ray structure of Qtud-3. **A**. *Left*, X-Ray structure of Q-eTud3 consisting of an extended Tudor fold, where the Tudor domain (magenta) separates the N-terminal portion of the SN domain (cyan) from the C-terminal one (blue). A long α-helical segment (yellow) connects the two domains. *Middle*, expanded view of the aromatic cage of Q-eTud3 (magenta) overlayed with the aromatic cage of P100 (blue, PDB ID 3BDL), which recognizes sDMA. *Right*, the electrostatic map of Q-eTud3 generated by the APBS module of Pymol showing the presence of a positively charged groove between the two domains. **B**. Comparison of the SN domains of Q-eTud3, Staphylococcal nuclease (PDB ID 2SNS) and P100 (PDB ID 3BDL); differently from the other SN domains, the SN domain of Q-eTud3 features a long β-strand-loop-β-strand segment extending from β9.

Another important difference between the SN domain of the Staphylococcal Nuclease and that of Q-eTud3 is the lack of the Ω-loop. Staphylococcal Nuclease has Ca^2+^-dependent DNase activity, which was found to depend on residue E43, located at the basis of the Ω-loop (amino acids 43–52) (Hibler et al., 1987); in addition, deletion of Ω-loop amino acids 44–49 increases activity (Poole et al., 1991). The Ω-loop is part of a 19-aa long linker extending between β-strand 7 and α-helix 3 (numbering according to Q-eTud3); in Q-eTud3 the β7–α3 linker is 6 aa shorter than in the SN domains of Staphylococcal Nuclease and the Ω-loop does not exist.

The Tudor domain has the typical four β-strands β-barrel topology, with a small helix located beside the barrel, and is intercalated between the second and third β-strands of the SN domain (β2 and β7 in Fig. 2A), with which it is connected via a long helical linker. In Tudor domains the aromatic cage is responsible for the recognition of methylated arginine and lysine marks (Tripsianes et al., 2011). This structural element is formed by aromatic residues located in the loops between the second and third as well as between the fourth and fifth β-strands of the Tudor domain (β3-β4 and β5-β6 in Fig. 2A). Overlaying the aromatic cage of P100 with the corresponding motif in Q-eTud3 shows that three out of the four aromatic residues building the cage in P100 are substituted by aliphatic residues in Q-eTud3 (Fig. 2A). The lack of a canonical aromatic cage explains the inability of Q-eTud3 to recognize methylated ligands.

To test if the other four Q-eTuds could adopt a similar fold, we generated high-confidence homology models of the Q-eTuds, which all show a similar extended Tudor topology. To verify the validity of the homology models, Q-eTud5 was subjected to small-angle X-ray scattering analysis; we obtained an excellent fit between the experimental data and the theoretical scattering curve generated from the homology model, confirming the predicted extended Tudor topology (Fig. S3F).

Most of the structural variability among the five Q-eTuds clusters around the SN domain (Fig. S3 and S4). Neither Q-eTud1 nor Q-eTud4 have the β-strand–loop–β-strand motif extending from β9, while this insertion is much shorter in Q-eTud2 than in Q-eTud3. In Q-eTud5 the β-strand–loop–β-strand motif extends from β8 rather than β9 and thus protrudes in the opposite direction.

Next, we inspected the region corresponding to the aromatic cage of the Tudor domain in all homology models. In both Q-eTud1 and Q-eTud2, all aromatic residues of the canonical aromatic cage are replaced by hydrophobic or charged residues, explaining why Q-eTud12 is unable to bind methylated ligands. In contrast, Q-eTud4 displays a canonical aromatic cage similar to that of TDRD3, which binds aDMA (Sikorsky et al., 2012). In particular, Y566, which was found to be crucial for the recognition of aDMA and for the discrimination between aDMA and sDMA in TDRD3, is conserved in Q-eTud4 (Y1344), in agreement with its ability to specifically recognize aDMA. Q-eTud5 shows an aromatic cage similar to that of P100; however, while P100 binds sDMA, Q-eTud5 was unable to bind any methylated ligand in our experiments. Two notable differences between the aromatic cage of the two proteins are an asparagine to aspartate (N743 to D1751) and a phenylalanine to tyrosine (F715 to Y1717) substitution (Fig. S3E). We do not know how these mutations may act to preclude sDMA binding.

In summary, while Q-eTud4 may bind aDMA marks, the function of Q-eTud1, Q-eTud2, Q-eTud3 and Q-eTud5 remains elusive.

### Qin’s extended Tudor domains are RNases

The SN fold originates from the Staphylococcal Nuclease protein, which shows endonuclease activity (Hynes and Fox, 1991; Li et al., 2018; Ponting, 2008). Thus, we tested whether the extended Tudor domains of Qin also have nuclease activity. For this purpose, we designed four structurally different RNAs and subjected them to cleavage assays with the four Qin constructs Qtud-12, Qtud-3, Qtud-34 and Qtud-5. The four RNA substrates were designed to adopt different secondary structures: single-stranded (ssRNA), double-stranded (dsRNA), stem-loop-stem (slsRNA) and stem-loop (slRNA) (Fig. 3 and Figs. S5 and S6). Both the monomeric and the dimeric form of Q-eTud3 (Q-eTud3m and Q-eTud3d, respectively) were tested.

**Figure 3.**
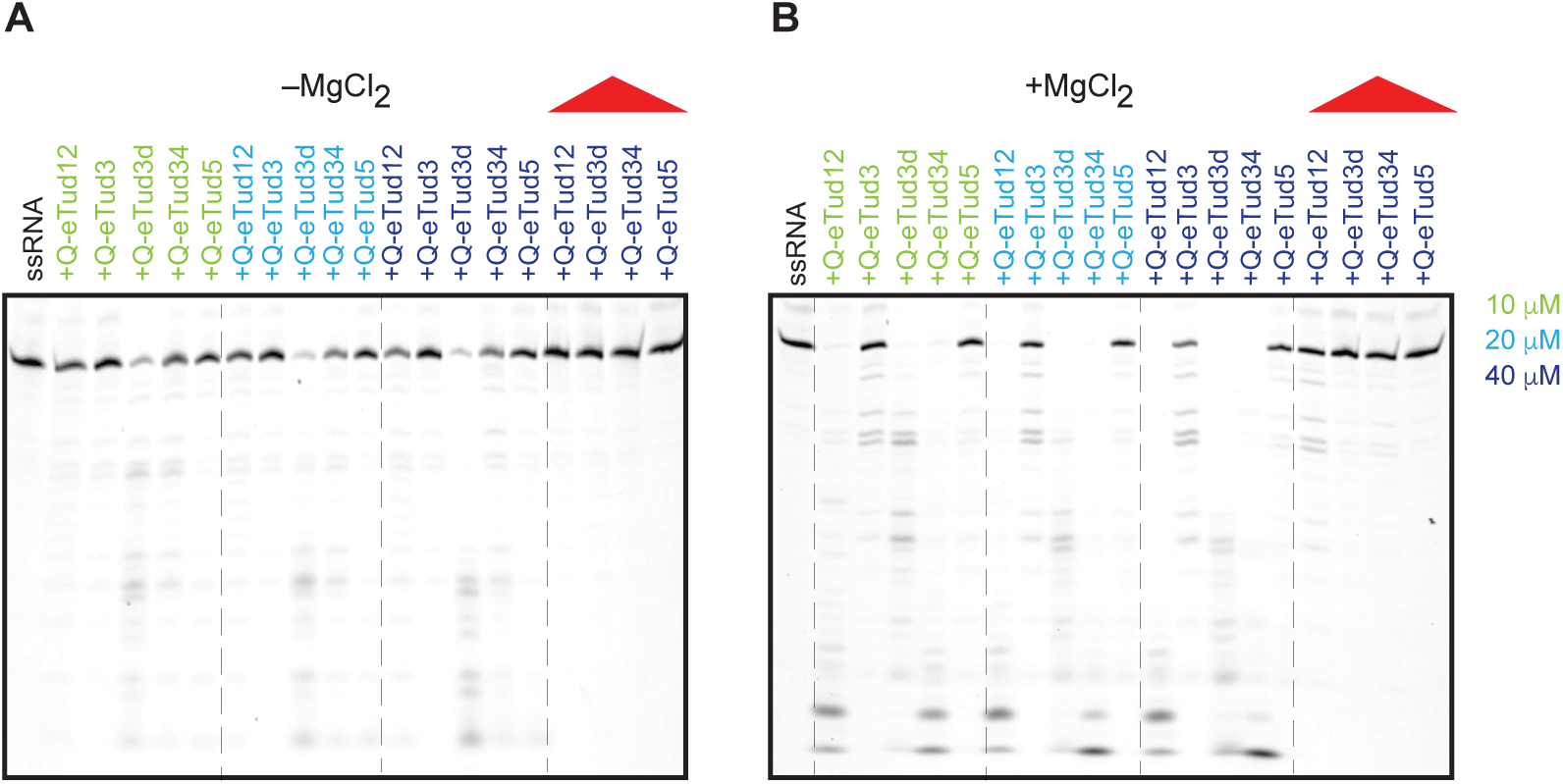
Q-eTuds act as RNases. Q-eTud12, Q-eTud3m (monomeric form), Q-eTud3d (dimeric form), Q-eTud34 and Q-eTud5 were assessed for RNase activity against ssRNA in buffer containing 25 mM HEPES potassium salt pH 7.0, 125 mM KCl, 1 mM DTT in the presence **(B)** and absence **(A)** of MgCl_2_ at 10 mM. The denaturing PAGE shows ∼0.5 μM of the 3’ end fluorescent labeled RNA after 5.5 hours of incubation at 30 °C with the five Q-eTud proteins. The concentrations of the Q-eTuds varied from 10 to 40 μM and are color coded according to the legend. The control lanes, containing RNA incubated with heat-denatured protein in buffer, are labeled with a red triangle. The dashed lines separate the lanes containing either different Q-eTuds concentrations or the heat-denatured proteins.

RNA digestion was monitored by the decrease in the intensity of the band corresponding to the full-size RNA as well as by the appearance of RNAs of smaller size. As controls, we used both the RNAs without any protein and heat-denatured solutions of the proteins; the last controls were necessary to rule out that the observed effects were due to contamination with RNases. In addition, with tested the proteins in the presence of either Mg^2+^ or EDTA to evaluate whether their activity is dependent on the presence of divalent metals.

All Q-eTud constructs were able to cleave ssRNA with the exception of Q-eTud5 (Figs. 3 and S5), which had negligible nuclease activity. None of the controls showed any nuclease activity, excluding that the RNA processing observed in the presence of the Q-eTud constructs was due to contamination with bacterial RNases. None of the Q-eTuds was active against dsRNA (Fig. S5) or stem-loop RNAs (Fig. S6), with the exception of the dimeric Q-eTud3d. The presence of 10 mM Mg^2+^ stimulated the RNase activity all constructs with the exception of Q-eTud5, which remained inactive (Figs. 3 and S5). RNase assays with increasing concentrations of Q-eTuds in the presence and absence of Mg^2+^ showed that 10 μM of either Q-eTud12 or Q-eTud34 in the presence of Mg^2+^ cleaved 0.5 μM ssRNA better than 40 μM protein in the absence of the cation (Fig. 3). Thus, we concluded that the activity of Q-eTud12 and Q-eTud34 increases by more than 4-fold upon addition of Mg^2+^.

The dimeric Q-eTud3d cleaved ssRNA more efficiently than the corresponding monomer and, similarly to all other active constructs, its activity was enhanced by the presence of Mg^2+^ (Fig. 3). However, unlike all other constructs, Q-eTud3d cleaved also RNAs with secondary structure (Figs. S5 and S6). Although we tested Q-eTud3d in all our assays, as we were intrigued by its high activity, we don’t believe that this form has any significance in a cellular context.

Finally, because the Staphylococcal Nuclease is an unspecific nuclease, we tested Q-eTud1–4 constructs for DNase activity. The DNAs were designed as ssDNA and dsDNA using the same fluorescent labeling scheme as for the RNAs. None of the Q-eTud constructs revealed any measurable DNase activity (Fig. S7), confirming that Qin is a specific RNA nuclease.

### Q-eTuds act as endoribonucleases and produce 5’-monophosphate products

To understand the function of Qin during the ping-pong cycle, we tested whether its Tudor domains act as exo- or endoribonucleases. Both functions are required in the piRNA biogenesis pathways, with Argonaute proteins and Zucchini acting as endonucleases (Cenik & Zamore, 2011; Ipsaro, Haase, Knott, Joshua-Tor, & Hannon, 2012; Nishimasu et al., 2012; W. Wang et al., 2015), and Nibbler acting as an exonuclease (Han, Hung, Weng, Zamore, & Ameres, 2011; N. Liu et al., 2011; H. Wang et al., 2016). For Qin, 3’→ 5’ exonuclease activity could be ruled out, due to the presence of the bulky cyanin-5 dye on the 3’ end of the RNAs used as substrates in the cleavage assays. We probed 5’→ 3’ exonuclease activity by blocking access to the 5’ end of the ssRNA substrate through a 7-methylguanylate cap (m7G cap). The RNase assays were then performed with both capped and uncapped RNA, both of which were also labeled with cyanin-5 at the 3’ end. Both capped and uncapped ssRNAs were cleaved by Q-eTud1–4, demonstrating that the 5’-m7G cap does not interfere with the nuclease activity (Fig. 4A). Thus, we conclude that the Qin extended Tudor domains are endoribonucleases.

**Figure 4.**
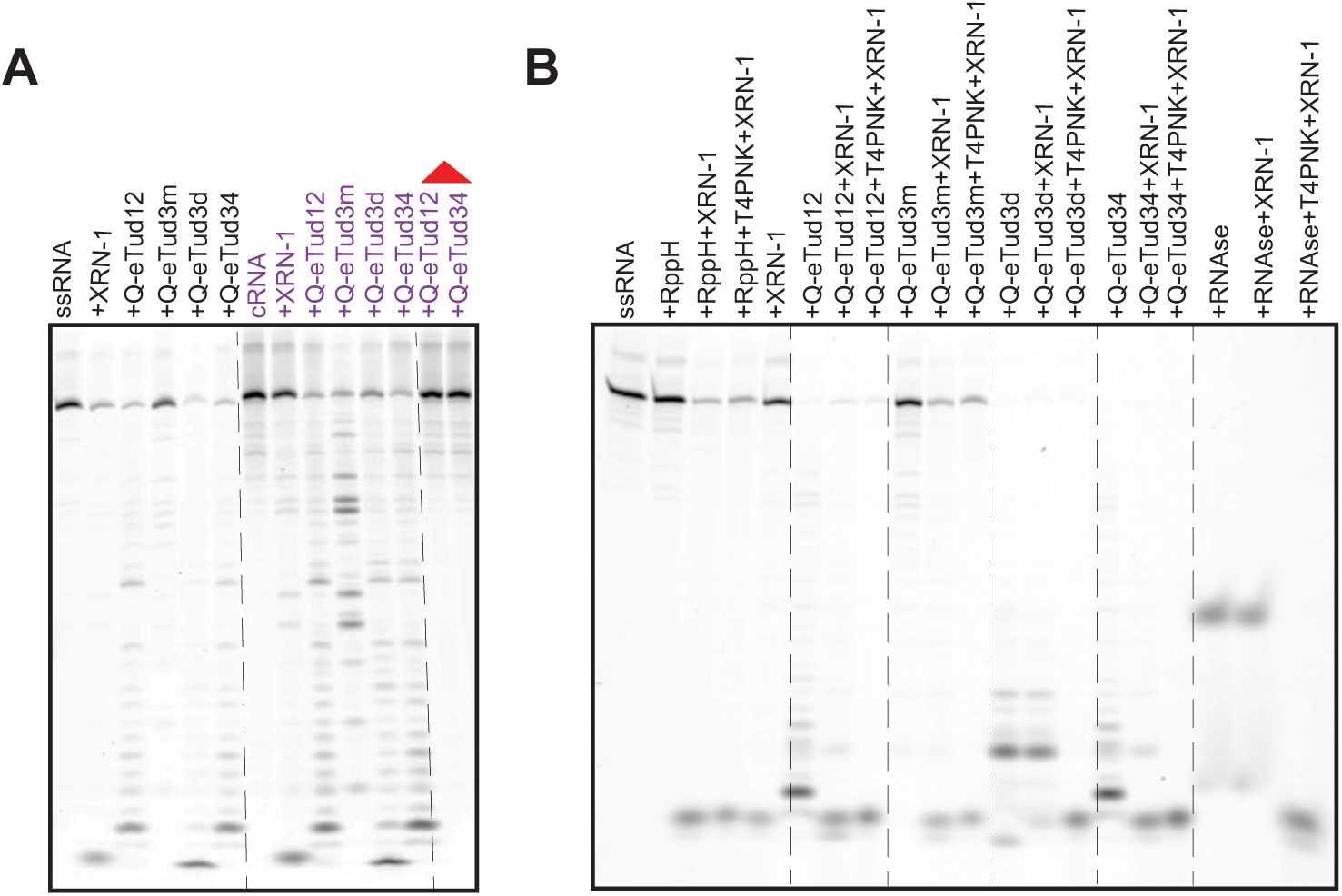
Q-eTuds are endonucleases and generate 5’-monophosphate ends. **A**. The RNase activity of Q-eTuds was evaluated upon incubation for 5.5 hours at 30 °C of ∼0.5 μM 3’ fluorescently-labeled ssRNA with (cRNA, purple) and without (ssRNA, black) a 5’-7mG cap together with 40 μM of Q-eTud (buffer composition: 10 mM MgCl_2_, 25 mM HEPES potassium salt pH 7.0, 125 mM KCl, 1 mM DTT). The 5’-cap does not inhibit cleavage of the substrate RNA. The control lanes, containing RNA incubated with heat-denatured protein in the same buffer, are labeled with a red triangle. The dashed lines separate the lanes containing either capped or uncapped ssRNA or the heat-denatured proteins **B**. The 5’-monophosphate specific 5’→3’ exonuclease XRN-1 and the T4 polynucleotide kinase T4PNK (which phosphorylates the RNA 5’-OH ends) were used to probe the nature of the 5’ ends of the RNA cleavage products generated by the Q-eTud constructs. XRN-1 was capable of digesting the cleavage products of all Q-eTud constructs without prior application of T4PNK, with the exception of Q-eTud3d. In all nuclease assays, 40 μM of Q-eTud proteins were incubated with 0.5 μM of 3’ end fluorescent labeled RNA for 5.5 hours at 30 °C in buffer containing 25 mM HEPES potassium salt pH 7.0, 125 mM KCl, 1 mM DTT and 10 mM MgCl_2_. The proteins were heat-inactivated before the addition of XRN-1 or T4PNK. The RNA-only control was treated with RNA 5’ pyrophosphohydrolase (RppH) before addition of XRN-1 or T4PNK. The dashed lines separate the lanes containing different Q-eTud proteins.

To test which chemical groups are left behind by the nucleolytic reaction catalysed by the Q-eTuds, we subjected the products of the nuclease assays to digestion with the 5’→3’ exonuclease XRN-1 that specifically recognizes 5’-monophosphate. After incubation with the Q-eTuds, the proteins were heat-inactivated and the RNA products were either treated directly with XRN-1 or pretreated with T4 polynucleotide kinase (T4PNK), which reestablishes the phosphate group on 5’-OH RNA ends, before addition of the exonuclease. XRN-1 was able to digest the cleavage products of Q-eTud12, Q-eTud3m and Q-eTud34 without pretreatment with T4PNK and the degradation pattern upon addition of either XRN-1 alone or first T4PNK and then XRN-1 remained identical. By contrast, the products of dimeric Q-eTud3d and the control RNaseA required pretreatment with the kinase to be further digested by XRN-1 (Fig. 4B). We conclude that all Q-eTud domains cleave the RNA yielding 5’-monophosphate and 3’-OH ends, with the exception of Q-eTud3d.

### Residues in the SN fold of Q-eTud3 and Q-eTud4 are important for nuclease activity

Last we probed whether the endonuclease activity of Qin’s extended Tudor domains depends on the SN fold. For this purpose, we inspected the structure of Q-eTud3 and compared it with the structure of the SN protein (Hynes & Fox, 1991). In the SN protein, mutations D21E/Y, R35K/G, D40E/G, E43D and R87K/G were proven detrimental for activity (Hale, Poole, & Gerlt, 1993; Pourmotabbed, Dell’Acqua, Gerlt, Stanczyk, & Bolton, 1990) (Serpersu, Shortle, & Mildvan, 1987), while Y113 and Y115 were shown to contact the bound nucleotide (Hynes & Fox, 1991) (Fig. 5A). Superposition of the SN structure with that of Q-eTud3 showed that Q-eTud3 D990, H1104, R1109 and E1114 occupy similar positions as SN D21, R35, D40 and E43 (Fig. 5A and B). Thus, we designed mutants D990A, H1104A/H1106A, R1109A and D1113A/E1114A and probed their activity as RNases. In addition, we generated mutant R1109A/E1181A/D1183A to test the role of negatively charged side chains in the vicinity of R1109A. We could produce all mutants with the exception of D990A, which was poorly expressed in *E. coli*. The H1104A/H1106A and R1109A mutations increased the nuclease activity of both Q-eTud3m and Q-eTud3d, while the R1109A/E1181A/D1183A and D1113A/E1114A mutants were either as active as or slightly less active than wild-type Q-eTud3m and Q-eTud3d (Fig. 5D).

**Figure 5.**
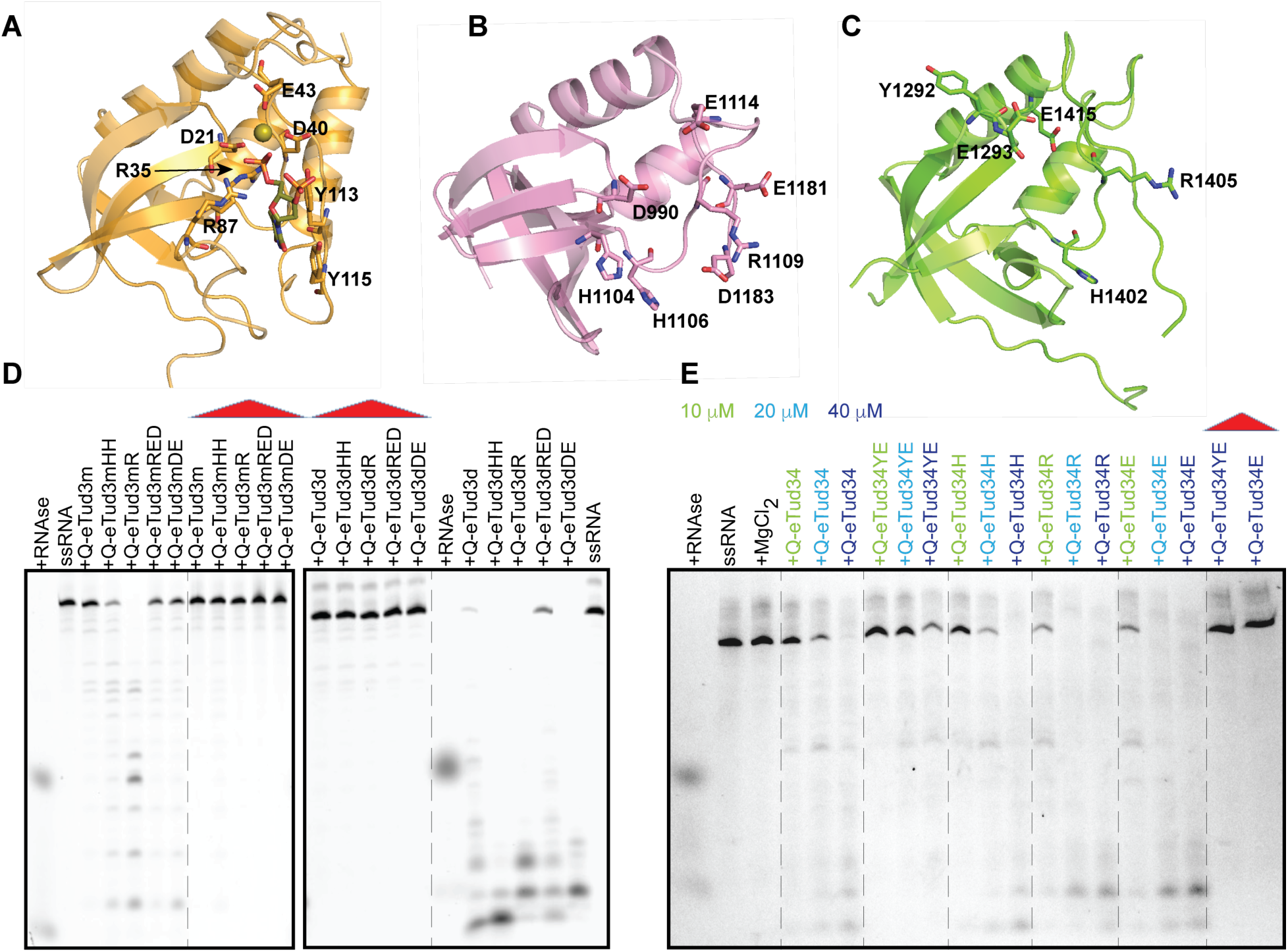
The nuclease activity of Qtud-5 resides in the SN fold. **A**. Structure of Staphylococcus Nuclease bound to the competitive inhibitor thymidine 3’,5’-bisphosphate and Ca^2+^ from PDB entry 2SNS (protein, light orange; ligand, olive; calcium ion, olive sphere). The amino acids important for either catalysis or substrate binding are shown in sticks. **B**. Structure of the SN domain of Q-eTud3 determined here. The amino acids shown in sticks occupy similar position as the catalysis relevant residues of SN and were mutated in nuclease assays in D to probe their relevance for activity. **C**. Homology model of the SN domain of Q-eTud4. The amino acids shown in sticks occupy similar position as the catalysis relevant residues of SN and were mutated in the nuclease assays in E to probe their relevance for activity. **D**. RNase assay of Q-eTud3 mutants. Q-eTud3m and Q-eTud3d indicate the monomeric and dimeric forms of the protein, respectively. The mutants tested were Q-eTud3 H1104A/H1106A (Q-eTud3HH), Q-eTud3 R1109A (Q-eTud3R) and Q-eTud3 D1113A/E1114A (Q-eTud3DE) and Q-eTud3 R1109A/E1181A/D1183A (Q-eTud3RED). The RNase activity of the Q-eTuds was evaluated upon incubation for 5.5 hours at 30 °C of ∼0.5 μM 3’ fluorescently-labeled ssRNA with 40 μM Q-eTud in the presence of 10 mM MgCl_2_ (buffer composition: 10 mM MgCl_2_, 25 mM HEPES potassium salt pH 7.0, 125 mM KCl, 1 mM DTT). The dashed lines separate the lanes containing heat-denatured proteins. **E**. RNase assay of Q-eTud4 mutants in the Q-eTud34 construct. The mutants tested were Q-eTud34 Y1292A/E1293A (Q-eTud34YE), Q-eTud34 H1402A (Q-eTud34H), Q-eTud34 R1405A (Q-eTud34R) and Q-eTud34 E1415A (Q-eTud34E). The RNase activity of the Q-eTuds was evaluated upon incubation for 5.5 hours at 30 °C of ∼0.5 μM 3’ fluorescently-labeled ssRNA with 10, 20 or 40 μM Q-eTud34 in the presence of 10 mM MgCl_2_ (buffer composition: 10 mM MgCl_2_, 25 mM HEPES potassium salt pH 7.0, 125 mM KCl, 1 mM DTT). The protein concentrations are color-coded according to the legend. The control lanes, containing RNA incubated with heat-denatured protein in the same buffer, are labeled with a red triangle. The dashed lines separate the lanes containing either different Q-eTud proteins or heat-denatured proteins.

We followed the same criteria to design mutations of the Q-eTud4 domain in the Q-eTud34 construct. The conservation of critical SN residues in Q-eTud4 is poor. As in Q-eTud3, an arginine (R1405) occupies the position of SN D40, while no charged residue is found at the positions corresponding to D21 and E43. Thus, we choose to mutate two glutamates whose sidechains are located in the same pocket as that of E43 in staphylococcal SN, namely E1293 and E1415. In addition, we mutated H1402, which lines up the pocket occupied by the nucleotide in staphylococcal SN (Fig. 5C). In total we tested the activity of four mutant Q-eTud34: Y1292A/E1293A, H1402A, R1405A and E1415A (Fig. 5E). The Y1292A/E1293A mutation decreased the activity of Q-eTud34, while the mutants R1405A and E1415A were more active than wild-type Q-eTud34. Finally, the H1402A mutation did not have any effect on activity. Interestingly, while the Q-eTud34 R1405A was more active than wild-type Q-eTud34 in the presence of magnesium, it was less active in its absence, suggesting that R1405 is detrimental for the stimulation of activity through the divalent cation (Fig. S8). An increase in nuclease activity in the presence of magnesium was measured also for the corresponding R1109A mutation in Q-eTud3, while depletion of the negative charges around R1109 in the R1109A/E1181A/D1183A neutralized the activity gain of the R1109A mutant (Fig. 5D). Taken together, these data suggest that the Mg^2+^ binding site is located in proximity of R1109 and R1405 in Q-eTud3 and Q-eTud4, respectively.

In conclusion, our results confirm that the SN domain harbors important residues for the nuclease activity of both Q-eTud3 and Q-eTud4; however, as the mutation of the Q-eTud3 glutamic acid corresponding to staphylococcal SN E43 did not impact activity, and Q-eTud4 lacks this glutamic acid in its native sequence, the enzymatic mechanism of Q-eTud34 should differ substantially from that of staphylococcal SN.

## DISCUSSION

The five C-terminal domains of Qin/Kimo adopt an extended Tudor fold, as other Tudor domains contained in the piRNA processing pathway. Q-eTud1–3 lack the aromatic cage motif and fail to recognize any methylated amino acid mark. Of the two Q-eTuds with an intact aromatic cage, Q-eTud5 does not bind any methylated ligands *in vitro*, while Q-eTud4 recognizes aDMA marks at high concentration. The affinity of the interaction remains weak when the aDMA motif is embedded in peptide sequences from Vasa, the only member of the amplification complex known to contain the aDMA modification (Kirino et al., 2010), thus questioning the significance of the Q-eTud4–aDMA interaction in this context. The atypical behavior of the Q-eTuds adds to the peculiar functions of other extended Tudor domains in the piRNA pathway: despite the conservation of their aromatic cages, the extended Tudor domains of Papi and Krimper recognize both methylated and non-methylated peptides (Webster et al., 2015; Y. Zhang et al., 2018), while Tdrd2, which lacks an aromatic cage, recognize non-methylated peptides only (H. Zhang et al., 2017). Altogether, these findings suggest that extended Tudor domains of the piRNA pathway function beyond the recognition of methylated ligands.

To explore additional functions of Q-eTuds, we considered the portion of the fold consisting of the SN domain. SN domains derive from the 149 amino-acid-long Staphylococcal Nuclease, a potent but non-specific Ca^2+^-dependent nuclease. The human homolog of SN (Ponting, 1997), protein P100, containing four sequential SN domains and an extended Tudor domain, cleaves ssRNA and binds dsRNA. The Ca^2+^-dependent nuclease function is harbored by the four SN domains, while the extended Tudor domain mediates the interaction with snRNPs or Piwi (Chia Lung Li, Yang, Shi, & Yuan, 2018) (Gao et al., 2012) (Shaw et al., 2007) and is involved neither in RNA binding nor in the nuclease activity at the tested concentration of 5 μM for 1 pmole RNA (C. L. Li et al., 2008). In contrast, we find that Q-eTud1–4 are Mg^2+^-dependent endonucleases, which preferentially cleave ssRNA to generate smaller RNAs with a 5’-phosphate group. Contrarily, Q-eTud5 does not have any nuclease activity and its function may involve binding to unmodified peptide stretches, as readily observed for Tdrd2 (H. Zhang et al., 2017).

The nuclease activity of tandem constructs, such as Q-eTud12 and Q-eTud34 is higher than that of isolated domains. This can either be due to a mere additive effect or involve cooperativity. Two Q-eTud domains can act cooperatively either by binding the RNA with stronger affinity, for example building a larger positively charged surface, or by cleaving the RNA more efficiently. Due to the unavailability of the single domains Q-eTud1, Q-eTud2 or Q-eTud4, we were unable to test the two hypotheses. In favor of the existence of cooperativity is the observation that a small percentage of isolated Q-eTud3 dimerizes and that Q-eTud3d is much more active than Q-eTud3m (Fig. 4). Q-eTud3d has different properties from the other Q-eTud constructs (for example, it cleaves also structured RNAs and its products have a 5’-OH rather than a 5’-monophosphate) and there is no evidence that these dimers are functional *in vivo*. However, the much higher *in vitro* activity of Q-eTud3d as compared with Q-eTud3m suggests that the Q-eTuds are optimized to act as multimers.

On the basis of our results, we propose that the function of Qin in piRNA biogenesis is linked to its RNase activity. During the ping-pong cycle, the splicer activity of Aub and Ago3 results in the formation of the 5’ end of the piRNA, while the 3’ end needs to be processed by ancillary proteins (Hayashi et al., 2016; Mohn, Handler, & Brennecke, 2015). Besides PIWI proteins, Squash, Zucchini, and Nibbler are the other known nucleases in the pathway. Squash has been shown to associate with Aub (Pane, Wehr, & Schupbach, 2007) but its mechanism of action remains elusive. Zucchini belongs to the superfamily of phospholipases-D/nucleases containing an HKD motif and falls in the sub-group of nucleases anchored to the mitochondrial membrane (Ipsaro et al., 2012; Pane et al., 2007). In primary piRNA biogenesis, Zucchini cleaves the cluster transcript with a 5’ U bias; in secondary piRNA biogenesis, Zucchini processes PIWI-bound pre-piRNAs to generate a Piwi-bound phased piRNA complex, which is then fed into the primary biogenesis pathway (X. Huang, Fejes Toth, & Aravin, 2017; Mohn et al., 2015) (Fig. 6). As a result, in the absence of Zucchini, Piwi-bound piRNAs are severely compromised. Conversely, the role of Zucchini in the ping-pong cycle is less obvious, as Zucchini mutants affect neither the ping-pong cycle (Han, Wang, Li, Weng, & Zamore, 2015) nor the generation of secondary piRNAs (Mohn et al., 2015). In the absence of Zucchini, the downstream (3’ end) U-bias of Aub-bound secondary piRNAs is diminished and the length of these piRNA is altered (Mohn et al., 2015), which indicates a role for Zucchini in 3’ end processing of secondary piRNAs loaded on Aub. Conversely, piRNAs associated with Ago3 are much less affected by Zucchini deletion, suggesting the existence of another nuclease that processes their 3’ end.

**Figure 6.**
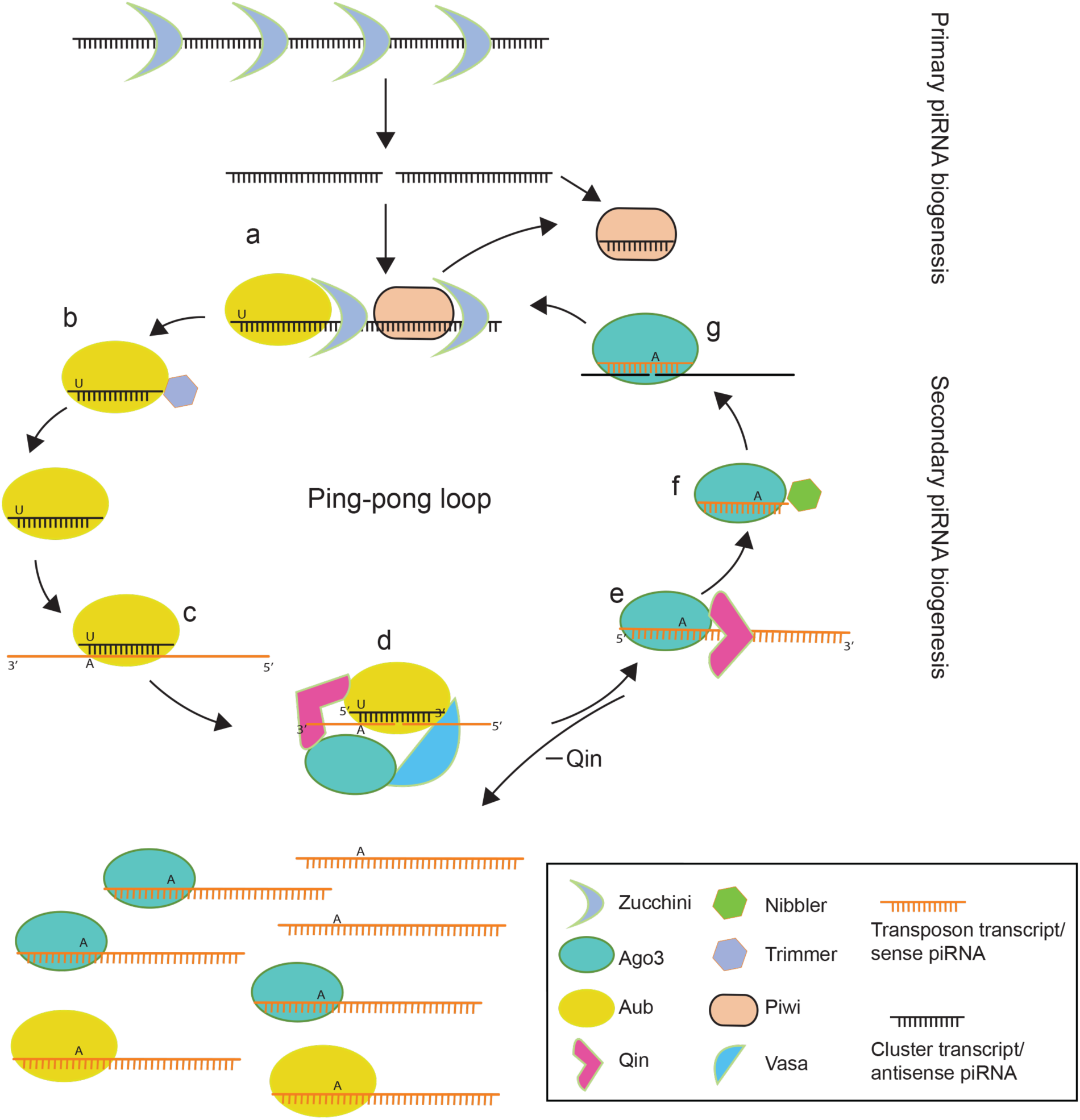
A model for the role of Qin in the ping-pong cycle. Scheme of the primary and secondary piRNA biogenesis processes with the proteins most relevant for RNA cleavage. Steps a and b of the ping-pong loop represent the processing of the 3’ end of antisense pre-piRNA bound to Aub; in c, the antisense pre-piRNA binds its complementary sequence in the transposon transcript; in d, the 5’ end of the sense pre-piRNA is generated by cleavage through Aub in the amplification complex; e and f represent the processing of the 3’ end of sense pre-piRNA bound to Ago3, according to the newly proposed mechanism, based on cleavage through Qin and trimming through Nibbler; in g, the sense piRNA recognizes the cluster transcripts and Ago3 cleavage generates the 5’ end of the antisense piRNA, which is then loaded onto Aub to reinitiate the cycle. In the absence of Qin, an excess of sense pre-piRNA is produced and loaded also on Aub, where it can be processed by Zucchini (as in a and b) to generate an Aub–sense piRNA complex (homotypic ping-pong).

Nibbler is the third nuclease acting in piRNA biogenesis. This 3’→5’ exonuclease has been proposed to trim the 3’ end of piRNAs prior to methylation by Hen1 (X. Huang et al., 2017; Kawaoka, Izumi, Katsuma, & Tomari, 2011). In the absence of Nibbler, piRNAs accumulate that are ∼0.6 nucleotide longer at the 3’ end (H. Wang et al., 2016), leading to the hypothesis that Nibbler fine-tunes the length of piRNAs. In contrast Hayashi et al. suggested that Zucchini and Nibbler act in two parallel pathways, with Zucchini processing the 3’ ends of most Aub-bound pre-piRNAs and Nibbler processing the 3’-end of most Ago3-bound pre-piRNAs (Hayashi et al., 2016). Nevertheless, piRNA mapping to many transposable elements is not affected by depletion of both Nibbler and Zucchini and the total level of piRNA in the germline is reduced by less than two-fold in comparison to Zucchini only depleted ovaries (Hayashi et al., 2016). This fact indicates the presence of another 3’ end processing enzyme that acts in parallel to Zucchini and Nibbler in the ping-pong cycle. Because Nibbler is absent in many mammal species, including rodents, this nuclease may be the dominant enzyme processing Ago3-loaded pre-piRNAs in these species.

In contrast to the loss of either Zucchini and Nibbler, disruption of Qin has remarkable consequences on the ping-pong cycle. Either knockout of Qin or deletion of its five extended Tudor domains results in an increase of the sense piRNA fraction for all families of sequenced transposable elements, while the antisense piRNA bias is lost because of a non-functional heterotypic ping-pong cycle (Z. Zhang et al., 2011). Instead, a homotypic ping-pong occurs between Aub:Aub, where the sense and antisense piRNA both bind to Aub. The same phenotype is observed upon Ago3 deletion (Senti, Jurczak, Sachidanandam, & Brennecke, 2015; Z. Zhang et al., 2011). In addition, the transposon families showing an increase in the transcript level upon Qin mutation are all known to require Ago3 for silencing, reinforcing the hypothesis of a functional correlation between Qin and Ago3 (Z. Zhang et al., 2011).

Based on our results and on the similarity of the phenotypes for the Qin and Ago3 functional mutants, we propose that Qin is the main endonuclease responsible for the maturation of the sense piRNAs bound to Ago3. In our new model of the ping-pong cycle (Fig. 6), the absence of Qin causes the heterotypic ping-pong to halt after the loading of the sense piRNAs on Ago3. Failure to process the 3’ end to yield mature sense piRNAs results in Ago3 stalling and leads to the same phenotype as Ago3 depletion. The excess of sense piRNAs, resulting from the lack of 3’ end processing on Ago3, may rebind to Aub, yielding a futile ping-pong cycle (Fig. 6). In support of this model, in Qin mutant ovaries, the excess sense piRNA fraction, with a A bias at position 10, was found to be loaded on Aub (Zhang et al., 2011). This model is not in contrast with that proposed by Hayashi et al. (Hayashi et al., 2016): Qin-dependent endonucleolytic cleavage could occur before exonucleolytic trimming by Nibbler and could account for the conservation of the ping-pong cycle in Zucchini and Nibbler depletion mutants.

Finally, the nuclease activity of Q-eTud1–4 measured in our assays *in vitro* is modest and is enhanced by the concerted action of multiple Q-eTuds. High nuclease activity of Q-eTuds against naked RNA, as tested in our assays, would lead to random degradation of piRNA intermediates; instead, Qin-dependent RNA-cleavage needs to be localized and its efficiency regulated in a position-dependent manner. A deeper understanding of the mechanism of this process will have to await the availability of the three-dimensional structure of the amplifier complex, containing Aub, Ago3, Vasa, Qin and the RNA.

## METHODS

### Lead Contact and Materials Availability

Requests for further information, resources and reagents should be directed to and will be fulfilled by the Lead Contact, Teresa Carlomagno. Atomic coordinates and structure factors have been deposited in the Protein Data Bank (PDB: 7OC3).

### Protein Expression and Purification

Expression plasmids contained cDNA sequences from *Drosophila melanogaster*, codon optimized for expression in *E. coli* (UniProtKb accession code Q9VE55). The DNA sequences were amplified and cloned into pETM11 (Q-eTud3 in NcoI and BamHI sites) or pETM11-Sumo3 expression vectors (Q-eTud12 and Q-eTud5 in BamHI and XhoI sites; and Q-eTud34 in BamHI and HindIII sites). The Quikchange® II XL Site-Directed Mutagenesis protocol was used to generate Q-eTud3 and Q-eTud34 mutants. All constructs were verified by DNA sequencing. Plasmids for Q-eTud3 and Q-eTud5 were transformed into BL21-CodonPlus (DE3)-RIL *E. coli* cells, while Q-eTud34 and Q-eTud12 were transformed into ArcticExpress (DE3) *E. coli* cells (both cell lines from Agilent). The expression of Q-eTud3 and Q-eTud5 was induced at A600 = 0.8 with 0.7 mM IPTG (isopropyl β-D-1-thiogalactopyranoside) for 20 hours at 16 °C. For Q-eTud12 and Q-eTud34, the expression was induced at A600 = 0.8 with 0.2 mM IPTG for 20 hours at 11 °C. The cells were grown in either LB medium or M9 minimal medium. Cells were harvested by centrifugation at 5000 g, and stored at –20°C. For NMR measurements, expression of sparsely deuterated,^15^N-labeled Q-eTud34 was achieved by growing cells in D2O based M9 minimal media containing 1 g/L ^15^NH4Cl as the sole nitrogen source. Multiple rounds (25%, 33%, 56%, 70% and 99%) of D2O adaptation were necessary for high-yield expression.

The cells were lysed by sonication in lysis buffer (25 mM Tris pH 7.8, 150 mM KCl, 5% glycerol, 10 mM imidazole, 5 mM β-mercaptoethanol) with 2 mM phenylmethylsulfonylfluorid (PMSF) or EDTA-free protease inhibitor cocktail tablet (Roche), 20 mg lysozyme (Sigma) and 30 units RQ1 RNase free DNase (Promega) per liter of culture. The lysate was clarified by centrifugation at 33,000 g and loaded on pre-equilibrated Ni-Sepharose 6 Fast Flow (HisTrap FF, GE Healthcare). The column was washed with 20-30 column volumes (CV) of lysis buffer, followed by 5-10 CVs of lysis buffer with the addition of 2 M LiCl, 500 mM NaCl and 350 mM KCl, and again with 5-10 CVs of lysis buffer. The washing with LiCl was repeated 2–3 times. The Q-eTuds were then eluted in 5 CVs of elution buffer (lysis buffer + 400 mM imidazole). Proteases SenP2 or TEV were added to the eluate, which was dialyzed against 60–100 X dialysis buffer (25 mM Tris-HCl pH 7.8, 150 mM KCl, 5 mM β-mercaptoethanol) at room temperature for 1–2 hours and then at 4 °C overnight. Q-eTud3 was purified further with reverse-His trap. The pooled peak fractions were further purified by anion exchange using a HiTrap Q HP column (GE Healthcare) in dialysis buffer. Q-eTud3 and Q-eTud5 were concentrated and loaded on a HiLoad Superdex 75 16/600 column (GE Healthcare) pre-equilibrated with size-exclusion buffer (25 mM Tris-HCl pH 7.8, 125 mM KCl, 1 mM DTT (Dithiothreitol)); Q-eTud12 and Q-eTud34 were concentrated and loaded on a Superdex 200 16/600 column (GE Healthcare) pre-equilibrated with size-exclusion buffer. The peak fractions were pooled and concentrated by Amicon® ultra-centrifugal filter units (EMD Millipore). The purity was assessed by SDS-PAGE and concentrations were determined using extinction coefficient obtained from the ProtParam tool (2005).

### NMR spectroscopy

NMR spectra were recorded at 293 K on Bruker Avance III HD 600 MHz and 850 MHz spectrometers, equipped with N2-cooled and He-cooled inverse HCN triple resonance cryogenic probeheads, respectively and running Topspin 3.2 software. NMR samples of ^15^N-labelled Q-eTud3 Q-eTud34 were prepared in NMR buffer (25 mM potassium phosphate pH 7.0, 125 mM KCl, 1 mM DTT (Dithiothreitol)) at a concentration of 400 μM and 50 μM respectively. The NMR spectra were processed with Topspin 3.2 (Bruker). Titration of ^15^N-labelled Q-eTud3 and Q-eTud34 (partially deuterated) with the aDMA were recorded as ^1^H-^15^N HSQC spectra (Bodenhausen & Ruben, 1980; Mori, Abeygunawardana, Johnson, & van Zijl, 1995).

### Thermal Shift Assay (TSA) screens

All the ligands (aDMA, sDMA, R, K, MML, DML, TML, m6A, 5mC and 7mG) were added to the NMR buffer at the final concentrations of 0.1 mM, 1 mM and 10 mM. For the peptides, final concentrations of 0.8 mM and 1.2 mM were used. Q-eTuds were added at final concentrations of 8 μM (Q-eTud3 and Q-eTud5), 2 μM (Q-eTud12) and 4 μM (Q-eTud34). For all experiments a control was set up with protein and ligand alone, dissolved in buffer and DMSO, respectively. All experiments were performed in duplicates. The experiment and data processing were performed as in (Kozak et al., 2016).

### Fluorescent labeling of RNA

The RNA sequences were cloned between EcoRI and PstI restriction sites in vector pUC19 and prepared by T7 directed in vitro transcription. RNAs were purified from urea-PAGE by the soak and crush method as described in (Petrov, Wu, Puglisi, & Puglisi, 2013). RNAs were fluorescently labeled with cyanine 5 at the 3’ end with the help of pCp-Cy5 (Jena Bioscience, 84179) and T4 RNA ligase I (NEB, M0204S). The reaction was conducted in the dark at 16 °C for 18 hs following the NEB protocol “3’ end labeling of RNA using T4 RNA Ligase 1”. The labeled RNAs were purified with the phenol:chloroform extraction method as described in (Sambrook & Russell, 2006) and precipitated with 70% ethanol + NaCl. The RNAs were re-suspended in water and stored at – 20°C.

RNA sequences:

ssRNA: 5’-ccacucccaccauuucacgcgcucccucaccucacauacuagauacaccaauacuuacuuuauaac-3’ dsRNA:

5’-ccauuacuauauuauaccaaguccgugcgugcccacacgcacggacuugguauaauauaguaauggcugc-3’ slsRNA: 5’-ggagugagguagugcuuggccgcuuccagcgugcugggggucuaaacuccgguacuaccucacuccc-3’ slRNA: 5’-gguaguuuggccgcagcgugcuucuaaacuccacuaccc-3’

DNA sequences (Merck-Sigma-Aldrich):

ssDNA: 5’-ccactcccaccatttcacgcgctccctcacctcacatactagatacaccaatacttactttataac-3’

dsDNA: 5’-ccattactatattataccaagtccgtgcgtgcccacacgcacggacttggtataatatagtaatggctgc-3’

### Nuclease activity assays

The RNAs were annealed (95 °C to 20 °C in ∼50 minutes) prior to use. The fluorescent RNAs (∼0.5 μM) were incubated with increasing concentrations of Q-eTuds (10 μM, 20 μM, 40 μM) to a final volume of 10 μL in buffer (25 mM HEPES potassium salt pH 7.0, 125 mM KCl, 1 mM DTT) at 30 °C for 5,5 hours. In the RNA-only control, the buffer replaced the protein. Further controls were done by incubating heat-denatured Q-eTuds (40 μM, 75 °C for 90 minutes) with RNA. To test the dependency of nuclease activity on magnesium, in some assays either 10 mM MgCl2 or 10 mM EDTA were added to the buffer, as indicated in the caption to each figure. The reaction was stopped by addition of 1 μL proteinase K (NEB, P8107S), followed by 15 min incubation at room temperature and addition of 20% formaldehyde). After centrifugation, the samples were run on 20% denaturing urea-PAGE in TBE buffer (89 mM Tris base, 89 mM boric acid, 2 mM EDTA pH 8.3). The gel fluorescence was imaged using a ChemiDoc™ MP Imaging System (Bio-Rad) (GE Healthcare). Similar experiments were performed for the DNase assays.

### Determination of cleavage product 5’ end

The RNase assays were conducted as described above. Afterwards, the samples containing Q-eTuds were incubated at 75 °C for 15 minutes to denaturate the Q-eTud nucleases. Control samples containing only RNA were treated with RNA 5’ Pyrophosphohydrolase (RppH, NEB, M0356S) to convert 5’-PPP RNA into 5’-P RNA. Afterwards all samples were incubated either with 0.5 units of the 5’ → 3’ exoribonuclease XRN-1, (NEB, M0338S) at 37 °C for 1 hour or first with 5 units T4 Polynucleotide Kinase with 25 mM ATP in 1 X T4 Kinase buffer (T4PNK, NEB, M0236S) at 37 °C for 30 min and then with XRN-1. Between the T4PNK treatment and the XRN-1 treatment, T4PNK was inactivated by incubation at 65 °C for 20 min. After treatment with proteinase K and formaldehyde followed by centrifugation, samples were subjected to gel electrophoresis on a 20% denaturing gel.

### Capping of RNA

Capping of RNA with the 5’-m7G cap was achieved using the Vaccinia Capping System (NEB), according to the manufacturer’s protocol. The RNA was then labeled with pCp-Cy5, as described above. After fluorescence labeling, the RNA was treated with 1 unit XRN-1 at 37 °C for 1,5 hours to remove any uncapped RNA. The RNA was purified with the NEB Monarch® RNA Cleanup Kit T2030S. The nuclease assays were performed as described above.

### Crystallization

Q-eTud3 was crystallized in the *P31*21 space group with two molecules per asymmetric unit and the structure was solved to a resolution of 2.3 Å with Se-methionine phasing (Table 1). Crystals of selenomethionine-labeled Q-eTud3 were grown at 4 °C by sitting-drop vapor diffusion. 0.150 μL of protein (14 mg/ml in NMR buffer with 1 mM TCEP (tris(2-carboxyethyl)phosphine)) was mixed with 0.150 μL of the crystallization solution (0.1 M Tris pH 8.5, 20% MPD). 20% (v/v) glycerol was used as the crystal cryo-protectant and was added before flash freezing the crystals in liquid nitrogen. Data collection and refinement statistics are given in Table 1. XDS, CCP4 suite, Refmac and Coot were used for data processing and refinement (Emsley, Lohkamp, Scott, & Cowtan, 2010; Kabsch, 2010; Murshudov et al., 2011; Winn et al., 2011).

### Small-angle X-ray Scattering (SAXS)

The data was collected at beamline P12 at the DESY light source Petra III at EMBL Hamburg (Blanchet et al., 2015). Q-eTud5 at a concentration of 8 mg/ml in 25 mM potassium phosphate buffer pH 6.0, 250 mM KCl, 5 mM DTT, was used for the SAXS experiments at 293 K. The frames were averaged and buffer subtracted, the inter-particle effects were removed and the curve was extrapolated to zero concentration using the ATSAS program suite. Crysol (Svergun, Barberato, & Koch, 1995) was used for the calculation of the theoretical scattering profile from the homology model of Q-eTud5 and its fitting to the experimental profile.

### Homology Modeling

Homology models were generated with the Protein Homology/analogY Recognition Engine V 2.0 (Phyre^2^) web portal (Kelley, Mezulis, Yates, Wass, & Sternberg, 2015).

## ACKNOWLEDGEMENTS

T.C. acknowledges financial support from the Deutsche Forschungsgemeinschaft (grant CA 294/14-1). We thank Dr. Melissa Ann Graewert for assistance in the collection of the SAXS data at the beam line P12 operated by EMBL Hamburg at the PETRA III storage ring (DESY, Hamburg, Germany). The crystallographic experiments were performed on beam line ID23 at the European Synchrotron Radiation Facility (ESRF), Grenoble, France. We are grateful to the Local Contact at the ESRF for providing assistance in using beam line ID23. We thank Dr. Stefan Schmelz at the Helmholtz Centre for Infection Research for collecting the MALS data. We acknowledge Life Science Editors for editing assistance.

## AUTHOR CONTRIBUTIONS

Conceptualization, T.C. and N.D.; Investigation, N.D., W.K. and S.L.; Writing –Original Draft, N.D. and T.C.; Writing –Review & Editing, N.D. and T.C.; Funding Acquisition, T.C.; Supervision, T.C.

## REFERENCES

Anand, A., & Kai, T. (2012). The tudor domain protein kumo is required to assemble the nuage and to generate germline piRNAs in Drosophila. EMBO J, 31(4), 870–882. doi:10.1038/emboj.2011.449

Andress, A., Bei, Y., Fonslow, B. R., Giri, R., Wu, Y., Yates, J. R., 3rd, & Carthew, R. W. (2016). Spindle-E cycling between nuage and cytoplasm is controlled by Qin and PIWI proteins. J Cell Biol, 213(2), 201–211. doi:10.1083/jcb.201411076

Aravin, A. A., Gaidatzis, D., Pfeffer, S., Lagos-Quintana, M., Landgraf, P., Iovino, N., … Tuschl, T. (2006). A novel class of small RNAs bind to MILI protein in mouse testes. Nature, 442(7099), 203–207. doi:10.1038/nature04916

Aravin, A. A., Hannon, G. J., & Brennecke, J. (2007). The Piwi-piRNA pathway provides an adaptive defense in the transposon arms race. Science, 318(5851), 761–764. doi:10.1126/science.1146484

Blanchet, C. E., Spilotros, A., Schwemmer, F., Graewert, M. A., Kikhney, A., Jeffries, C. M., … Svergun, D. I. (2015). Versatile sample environments and automation for biological solution X-ray scattering experiments at the P12 beamline (PETRA III, DESY). J Appl Crystallogr, 48(Pt 2), 431–443. doi:10.1107/S160057671500254X

Bodenhausen, G., & Ruben, D. J. (1980). Natural abundance nitrogen-15 NMR by enhanced heteronuclear spectroscopy. Chem. Phys. Lett., 69, 185–189.

Brennecke, J., Aravin, A. A., Stark, A., Dus, M., Kellis, M., Sachidanandam, R., & Hannon, G. J. (2007). Discrete small RNA-generating loci as master regulators of transposon activity in Drosophila. Cell, 128(6), 1089–1103. doi:10.1016/j.cell.2007.01.043

Cenik, E. S., & Zamore, P. D. (2011). Argonaute proteins. Curr Biol, 21(12), R446–449. doi:10.1016/j.cub.2011.05.020

Chen, C., Nott, T. J., Jin, J., & Pawson, T. (2011). Deciphering arginine methylation: Tudor tells the tale. Nat Rev Mol Cell Biol, 12(10), 629–642. doi:10.1038/nrm3185

Czech, B., & Hannon, G. J. (2016). One Loop to Rule Them All: The Ping-Pong Cycle and piRNA-Guided Silencing. Trends Biochem Sci, 41(4), 324–337. doi:10.1016/j.tibs.2015.12.008

Czech, B., Munafo, M., Ciabrelli, F., Eastwood, E. L., Fabry, M. H., Kneuss, E., & Hannon, G. J. (2018). piRNA-Guided Genome Defense: From Biogenesis to Silencing. Annu Rev Genet, 52, 131–157. doi:10.1146/annurev-genet-120417-031441

Elbarbary, R. A., Miyoshi, K., Myers, J. R., Du, P., Ashton, J. M., Tian, B., & Maquat, L. E. (2017). Tudor-SN-mediated endonucleolytic decay of human cell microRNAs promotes G1/S phase transition. Science, 356(6340), 859–862. doi:10.1126/science.aai9372

Emsley, P., Lohkamp, B., Scott, W. G., & Cowtan, K. (2010). Features and development of Coot. Acta Crystallogr D Biol Crystallogr, 66(Pt 4), 486–501. doi:10.1107/S0907444910007493

Gao, X., Zhao, X., Zhu, Y., He, J., Shao, J., Su, C., … Yang, J. (2012). Tudor staphylococcal nuclease (Tudor-SN) participates in small ribonucleoprotein (snRNP) assembly via interacting with symmetrically dimethylated Sm proteins. J Biol Chem, 287(22), 18130–18141. doi:10.1074/jbc.M111.311852

Girard, A., Sachidanandam, R., Hannon, G. J., & Carmell, M. A. (2006). A germline-specific class of small RNAs binds mammalian Piwi proteins. Nature, 442(7099), 199–202. doi:10.1038/nature04917

Grimson, A., Srivastava, M., Fahey, B., Woodcroft, B. J., Chiang, H. R., King, N., … Bartel, D. P. (2008). Early origins and evolution of microRNAs and Piwi-interacting RNAs in animals. Nature, 455(7217), 1193–1197. doi:10.1038/nature07415

Hale, S. P., Poole, L. B., & Gerlt, J. A. (1993). Mechanism of the reaction catalyzed by staphylococcal nuclease: identification of the rate-determining step. Biochemistry, 32(29), 7479–7487. doi:10.1021/bi00080a020

Han, B. W., Hung, J. H., Weng, Z., Zamore, P. D., & Ameres, S. L. (2011). The 3’-to-5’ exoribonuclease Nibbler shapes the 3’ ends of microRNAs bound to Drosophila Argonaute1. Curr Biol, 21(22), 1878–1887. doi:10.1016/j.cub.2011.09.034

Han, B. W., Wang, W., Li, C., Weng, Z., & Zamore, P. D. (2015). Noncoding RNA. piRNA-guided transposon cleavage initiates Zucchini-dependent, phased piRNA production. Science, 348(6236), 817–821. doi:10.1126/science.aaa1264

Hayashi, R., Schnabl, J., Handler, D., Mohn, F., Ameres, S. L., & Brennecke, J. (2016). Genetic and mechanistic diversity of piRNA 3’-end formation. Nature, 539(7630), 588–592. doi:10.1038/nature20162

Hibler, D. W., Stolowich, N. J., Reynolds, M. A., Gerlt, J. A., Wilde, J. A., & Bolton, P. H. (1987). Site-directed mutants of staphylococcal nuclease. Detection and localization by 1H NMR spectroscopy of conformational changes accompanying substitutions for glutamic acid-43. Biochemistry, 26(19), 6278–6286. doi:10.1021/bi00393a048

Huang, H. Y., Houwing, S., Kaaij, L. J., Meppelink, A., Redl, S., Gauci, S., … Ketting, R. F. (2011). Tdrd1 acts as a molecular scaffold for Piwi proteins and piRNA targets in zebrafish. EMBO J, 30(16), 3298–3308. doi:10.1038/emboj.2011.228

Huang, X., Fejes Toth, K., & Aravin, A. A. (2017). piRNA Biogenesis in Drosophila melanogaster. Trends Genet, 33(11), 882–894. doi:10.1016/j.tig.2017.09.002

Hynes, T. R., & Fox, R. O. (1991). The crystal structure of staphylococcal nuclease refined at 1.7 A resolution. Proteins, 10(2), 92–105. doi:10.1002/prot.340100203

Ipsaro, J. J., Haase, A. D., Knott, S. R., Joshua-Tor, L., & Hannon, G. J. (2012). The structural biochemistry of Zucchini implicates it as a nuclease in piRNA biogenesis. Nature, 491(7423), 279–283. doi:10.1038/nature11502

Kabsch, W. (2010). XDS. Acta crystallographica. Section D, Biological crystallography, 66(Pt 2), 125–132. doi:10.1107/S0907444909047337

Kawaoka, S., Izumi, N., Katsuma, S., & Tomari, Y. (2011). 3’ end formation of PIWI-interacting RNAs in vitro. Mol Cell, 43(6), 1015–1022. doi:10.1016/j.molcel.2011.07.029

Kelley, L. A., Mezulis, S., Yates, C. M., Wass, M. N., & Sternberg, M. J. E. (2015). The Phyre2 web portal for protein modeling, prediction and analysis. Nature Protocols, 10(6), 845–858. doi:10.1038/nprot.2015.053

Kirino, Y., Vourekas, A., Kim, N., de Lima Alves, F., Rappsilber, J., Klein, P. S., … Mourelatos, Z. (2010). Arginine methylation of vasa protein is conserved across phyla. The Journal of biological chemistry, 285(11), 8148–8154. doi:10.1074/jbc.M109.089821

Kozak, S., Lercher, L., Karanth, M. N., Meijers, R., Carlomagno, T., & Boivin, S. (2016). Optimization of protein samples for NMR using thermal shift assays. Journal of Biomolecular NMR, 64(4), 281–289. doi:10.1007/s10858-016-0027-z

Ku, H. Y., & Lin, H. (2014). PIWI proteins and their interactors in piRNA biogenesis, germline development and gene expression. Natl Sci Rev, 1(2), 205–218. doi:10.1093/nsr/nwu014

Lau, N. C., Seto, A. G., Kim, J., Kuramochi-Miyagawa, S., Nakano, T., Bartel, D. P., & Kingston, R. E. (2006). Characterization of the piRNA complex from rat testes. Science, 313(5785), 363–367. doi:10.1126/science.1130164

Lee, H.-E., Ayarpadikannan, S., & Kim, H.-S. (2015). Role of transposable elements in genomic rearrangement, evolution, gene regulation and epigenetics in primates. Genes & Genetic Systems, 90(5), 245–257. doi:10.1266/ggs.15-00016

Levin, H. L., & Moran, J. V. (2011). Dynamic interactions between transposable elements and their hosts. Nat Rev Genet, 12(9), 615–627. doi:10.1038/nrg3030

Li, C. L., Yang, W. Z., Chen, Y. P., & Yuan, H. S. (2008). Structural and functional insights into human Tudor-SN, a key component linking RNA interference and editing. Nucleic Acids Res, 36(11), 3579–3589. doi:10.1093/nar/gkn236

Li, C. L., Yang, W. Z., Shi, Z., & Yuan, H. S. (2018). Tudor staphylococcal nuclease is a structure-specific ribonuclease that degrades RNA at unstructured regions during microRNA decay. RNA (New York, N.Y.), 24(5), 739–748. doi:10.1261/rna.064501.117

Liu, H., Wang, J. Y., Huang, Y., Li, Z., Gong, W., Lehmann, R., & Xu, R. M. (2010). Structural basis for methylarginine-dependent recognition of Aubergine by Tudor. Genes Dev, 24(17), 1876–1881. doi:10.1101/gad.1956010

Liu, L., Qi, H., Wang, J., & Lin, H. (2011). PAPI, a novel TUDOR-domain protein, complexes with AGO3, ME31B and TRAL in the nuage to silence transposition. Development (Cambridge, England), 138(9), 1863–1873. doi:10.1242/dev.059287

Liu, N., Abe, M., Sabin, L. R., Hendriks, G. J., Naqvi, A. S., Yu, Z., … Bonini, N. M. (2011). The exoribonuclease Nibbler controls 3’ end processing of microRNAs in Drosophila. Curr Biol, 21(22), 1888–1893. doi:10.1016/j.cub.2011.10.006

Mathioudakis, N., Palencia, A., Kadlec, J., Round, A., Tripsianes, K., Sattler, M., … Cusack, S. (2012). The multiple Tudor domain-containing protein TDRD1 is a molecular scaffold for mouse Piwi proteins and piRNA biogenesis factors. RNA (New York, N.Y.), 18(11), 2056–2072. doi:10.1261/rna.034181.112

Mohn, F., Handler, D., & Brennecke, J. (2015). Noncoding RNA. piRNA-guided slicing specifies transcripts for Zucchini-dependent, phased piRNA biogenesis. Science (New York, N.Y.), 348(6236), 812–817. doi:10.1126/science.aaa1039

Mori, S., Abeygunawardana, C., Johnson, M. O., & van Zijl, P. C. (1995). Improved sensitivity of HSQC spectra of exchanging protons at short interscan delays using a new fast HSQC (FHSQC) detection scheme that avoids water saturation. J Magn Reson B, 108(1), 94–98. doi:10.1006/jmrb.1995.1109

Murshudov, G. N., Skubak, P., Lebedev, A. A., Pannu, N. S., Steiner, R. A., Nicholls, R. A., … Vagin, A. A. (2011). REFMAC5 for the refinement of macromolecular crystal structures. Acta Crystallographica Section D, 67(4), 355–367. doi:doi:10.1107/S0907444911001314

Nishida, K. M., Okada, T. N., Kawamura, T., Mituyama, T., Kawamura, Y., Inagaki, S., … Siomi, M. C. (2009). Functional involvement of Tudor and dPRMT5 in the piRNA processing pathway in Drosophila germlines. The EMBO journal, 28(24), 3820–3831. doi:10.1038/emboj.2009.365

Nishida, K. M., Sakakibara, K., Iwasaki, Y. W., Yamada, H., Murakami, R., Murota, Y., … Siomi, M. C. (2018). Hierarchical roles of mitochondrial Papi and Zucchini in Bombyx germline piRNA biogenesis. Nature, 555(7695), 260–264. doi:10.1038/nature25788

Nishimasu, H., Ishizu, H., Saito, K., Fukuhara, S., Kamatani, M. K., Bonnefond, L., … Nureki, O. (2012). Structure and function of Zucchini endoribonuclease in piRNA biogenesis. Nature, 491(7423), 284–287. doi:10.1038/nature11509

Ozata, D. M., Gainetdinov, I., Zoch, A., O’Carroll, D., & Zamore, P. D. (2019). PIWI-interacting RNAs: small RNAs with big functions. Nat Rev Genet, 20(2), 89–108. doi:10.1038/s41576-018-0073-3

Pan, J., Goodheart, M., Chuma, S., Nakatsuji, N., Page, D. C., & Wang, P. J. (2005). RNF17, a component of the mammalian germ cell nuage, is essential for spermiogenesis. Development, 132(18), 4029–4039. doi:10.1242/dev.02003

Pane, A., Wehr, K., & Schupbach, T. (2007). zucchini and squash encode two putative nucleases required for rasiRNA production in the Drosophila germline. Dev Cell, 12(6), 851–862. doi:10.1016/j.devcel.2007.03.022

Petrov, A., Wu, T., Puglisi, E. V., & Puglisi, J. D. (2013). RNA purification by preparative polyacrylamide gel electrophoresis. Methods Enzymol, 530, 315–330. doi:10.1016/B978-0-12-420037-1.00017-8

Ponting, C. P. (1997). P100, a transcriptional coactivator, is a human homologue of staphylococcal nuclease. Protein science : a publication of the Protein Society, 6(2), 459–463. doi:10.1002/pro.5560060224

Poole, L. B., Loveys, D. A., Hale, S. P., Gerlt, J. A., Stanczyk, S. M., & Bolton, P. H. (1991). Deletion of the .OMEGA.-loop in the active site of staphylococcal nuclease. I. Effect on catalysis and stability. Biochemistry, 30(15), 3621–3627. doi:10.1021/bi00229a005

Pourmotabbed, T., Dell’Acqua, M., Gerlt, J. A., Stanczyk, S. M., & Bolton, P. H. (1990). Kinetic and conformational effects of lysine substitutions for arginines 35 and 87 in the active site of staphylococcal nuclease. Biochemistry, 29(15), 3677–3683. doi:10.1021/bi00467a013

Ren, R., Liu, H., Wang, W., Wang, M., Yang, N., Dong, Y. H., … Xu, R. M. (2014). Structure and domain organization of Drosophila Tudor. Cell Res, 24(9), 1146–1149. doi:10.1038/cr.2014.63

Sambrook, J., & Russell, D. W. (2006). Purification of nucleic acids by extraction with phenol:chloroform. CSH Protoc, 2006(1). doi:10.1101/pdb.prot4455

Sato, K., Iwasaki, Y. W., Siomi, H., & Siomi, M. C. (2015). Tudor-domain containing proteins act to make the piRNA pathways more robust in Drosophila. Fly (Austin), 9(2), 86–90. doi:10.1080/19336934.2015.1128599

Senti, K. A., Jurczak, D., Sachidanandam, R., & Brennecke, J. (2015). piRNA-guided slicing of transposon transcripts enforces their transcriptional silencing via specifying the nuclear piRNA repertoire. Genes Dev, 29(16), 1747–1762. doi:10.1101/gad.267252.115

Serpersu, E. H., Shortle, D., & Mildvan, A. S. (1987). Kinetic and magnetic resonance studies of active-site mutants of staphylococcal nuclease: factors contributing to catalysis. Biochemistry, 26(5), 1289–1300. doi:10.1021/bi00379a014

Shaw, N., Zhao, M., Cheng, C., Xu, H., Saarikettu, J., Li, Y., … Rao, Z. (2007). The multifunctional human p100 protein ‘hooks’ methylated ligands. Nat Struct Mol Biol, 14(8), 779–784. doi:10.1038/nsmb1269

Sikorsky, T., Hobor, F., Krizanova, E., Pasulka, J., Kubicek, K., & Stefl, R. (2012). Recognition of asymmetrically dimethylated arginine by TDRD3. Nucleic Acids Res, 40(22), 11748–11755. doi:10.1093/nar/gks929

Siomi, M. C., Sato, K., Pezic, D., & Aravin, A. A. (2011). PIWI-interacting small RNAs: the vanguard of genome defence. Nat Rev Mol Cell Biol, 12(4), 246–258. doi:10.1038/nrm3089

Svergun, D., Barberato, C., & Koch, M. H. J. (1995). CRYSOL -a Program to Evaluate X-ray Solution Scattering of Biological Macromolecules from Atomic Coordinates. Journal of Applied Crystallography, 28(6), 768–773. doi:doi:10.1107/S0021889895007047

Tripsianes, K., Madl, T., Machyna, M., Fessas, D., Englbrecht, C., Fischer, U., … Sattler, M. (2011). Structural basis for dimethylarginine recognition by the Tudor domains of human SMN and SPF30 proteins. Nat Struct Mol Biol, 18(12), 1414–1420. doi:10.1038/nsmb.2185

Wang, H., Ma, Z., Niu, K., Xiao, Y., Wu, X., Pan, C., … Liu, N. (2016). Antagonistic roles of Nibbler and Hen1 in modulating piRNA 3’ ends in Drosophila. Development, 143(3), 530–539. doi:10.1242/dev.128116

Wang, W., Han, B. W., Tipping, C., Ge, D. T., Zhang, Z., Weng, Z., & Zamore, P. D. (2015). Slicing and Binding by Ago3 or Aub Trigger Piwi-Bound piRNA Production by Distinct Mechanisms. Mol Cell, 59(5), 819–830. doi:10.1016/j.molcel.2015.08.007

Webster, A., Li, S., Hur, J. K., Wachsmuth, M., Bois, J. S., Perkins, E. M., … Aravin, A. A. (2015). Aub and Ago3 Are Recruited to Nuage through Two Mechanisms to Form a Ping-Pong Complex Assembled by Krimper. Mol Cell, 59(4), 564–575. doi:10.1016/j.molcel.2015.07.017

Winn, M. D., Ballard, C. C., Cowtan, K. D., Dodson, E. J., Emsley, P., Evans, P. R., Wilson, K. S. (2011). Overview of the CCP4 suite and current developments. Acta Crystallogr D Biol Crystallogr, 67(Pt 4), 235–242. doi:10.1107/S0907444910045749

Xiol, J., Spinelli, P., Laussmann, M. A., Homolka, D., Yang, Z., Cora, E., Pillai, R. S. (2014). RNA clamping by Vasa assembles a piRNA amplifier complex on transposon transcripts. Cell, 157(7), 1698–1711. doi:10.1016/j.cell.2014.05.018

Zhang, H., Liu, K., Izumi, N., Huang, H., Ding, D., Ni, Z., … Min, J. (2017). Structural basis for arginine methylation-independent recognition of PIWIL1 by TDRD2. Proc Natl Acad Sci U S A, 114(47), 12483–12488. doi:10.1073/pnas.1711486114

Zhang, Y., Liu, W., Li, R., Gu, J., Wu, P., Peng, C., … Huang, Y. (2018). Structural insights into the sequence-specific recognition of Piwi by Drosophila Papi. Proc Natl Acad Sci U S A, 115(13), 3374–3379. doi:10.1073/pnas.1717116115

Zhang, Z., Koppetsch, B. S., Wang, J., Tipping, C., Weng, Z., Theurkauf, W. E., & Zamore, P. D. (2014). Antisense piRNA amplification, but not piRNA production or nuage assembly, requires the Tudor-domain protein Qin. EMBO J, 33(6), 536–539. doi:10.1002/embj.201384895

Zhang, Z., Xu, J., Koppetsch, B. S., Wang, J., Tipping, C., Ma, S., … Zamore, P. D. (2011). Heterotypic piRNA Ping-Pong requires qin, a protein with both E3 ligase and Tudor domains. Mol Cell, 44(4), 572–584. doi:10.1016/j.molcel.2011.10.011

