## Supplementary Figures 1-8 for "EXTENDED TUDOR-DOMAINS of the piRNA BIOGENESIS PATHWAY HAVE RNA-SPECIFIC NUCLEASE ACTIVITY"

### SUPPLEMENTAL INFORMATION

**Table S1: Binding of Q-eTud constructs to methylated and non-methylated ligands monitored by TSA.**

| | Q-eTud12 | | | Q-eTud3 | | | Q-eTud34 | | | Q-eTud5 | | |
|---|---|---|---|---|---|---|---|---|---|---|---|---|
| Ligand | [mM] | $T_m$ | S.D. | [mM] | $T_m$ | S.D. | [mM] | $T_m$ | S.D. | [mM] | $T_m$ | S.D. |
| C | - | 47,7 | 0,0 | - | 41,7 | 0,6 | - | 40,8 | 0,3 | - | 47,3 | 0,2 |
| sDMA | 0,1 | 47,4 | 0,0 | 0,1 | 41,4 | 0,0 | 0,1 | 40,8 | 0,0 | 0,1 | 47,3 | 0,2 |
| sDMA | 1 | 47,4 | 0,0 | 1 | 41,4 | 0,0 | 1 | 41,1 | 0,0 | 1 | 47,0 | 0,2 |
| sDMA | 10 | 47,6 | 0,2 | 10 | 41,3 | 0,2 | 10 | 41,3 | 0,2 | 10 | 47,3 | 0,2 |
| C | - | 47,7 | 0,0 | - | 41,7 | 0,6 | - | 40,2 | 0,0 | - | 47,1 | 0,3 |
| aDMA | 0,1 | 47,7 | 0,0 | 0,1 | 42,2 | 0,2 | **0,1** | **40,2** | **0,0** | 0,1 | 47,1 | 0,0 |
| aDMA | 1 | 47,7 | 0,0 | 1 | 41,7 | 0,0 | **1** | **40,5** | **0,0** | 1 | 47,1 | 0,0 |
| aDMA | 10 | 47,7 | 0,0 | 10 | 41,6 | 0,2 | **10** | **41,1** | **0,0** | 10 | 47,0 | 0,0 |
| Arg | 0,1 | 47,7 | 0,0 | 0,1 | 41,7 | 0,0 | 0,1 | 40,2 | 0,0 | 0,1 | 47,4 | 0,0 |
| Arg | 1 | 47,7 | 0,0 | 1 | 41,7 | 0,0 | 1 | 40,2 | 0,0 | 1 | 47,9 | 0,2 |
| Arg | 10 | 47,7 | 0,0 | 10 | 41,6 | 0,2 | 10 | 40,2 | 0,0 | 10 | 47,1 | 0,8 |
| Lys | 0,1 | 47,7 | 0,0 | 0,1 | 41,7 | 0,4 | 0,1 | 40,2 | 0,0 | 0,1 | 47,4 | 0,0 |
| Lys | 1 | 47,7 | 0,0 | 1 | 41,6 | 0,2 | 1 | 40,2 | 0,0 | 1 | 47,9 | 0,2 |
| Lys | 10 | 47,9 | 0,2 | 10 | 41,7 | 0,0 | 10 | 40,2 | 0,0 | 10 | 47,7 | 0,0 |
| MML | 0,1 | 47,7 | 0,0 | 0,1 | 41,9 | 0,2 | 0,1 | 40,2 | 0,0 | 0,1 | 47,6 | 0,2 |
| MML | 1 | 47,6 | 0,2 | 1 | 41,6 | 0,2 | 1 | 40,2 | 0,0 | 1 | 47,3 | 0,6 |
| MML | 10 | 47,6 | 0,2 | 10 | 41,4 | 0,4 | 10 | 40,2 | 0,0 | 10 | 47,4 | 0,4 |
| DML | 0,1 | 47,7 | 0,0 | 0,1 | 41,7 | 0,0 | 0,1 | 40,2 | 0,0 | 0,1 | 47,6 | 0,2 |
| DML | 1 | 47,6 | 0,2 | 1 | 41,4 | 0,4 | 1 | 40,2 | 0,0 | 1 | 47,4 | 0,0 |
| DML | 10 | 47,7 | 0,0 | 10 | 41,4 | 0,0 | 10 | 40,2 | 0,0 | 10 | 47,4 | 0,0 |
| TML | 0,1 | 47,6 | 0,2 | 0,1 | 40,8 | 2,1 | 0,1 | 40,2 | 0,0 | 0,1 | 47,6 | 0,2 |
| TML | 1 | 47,6 | 0,2 | 1 | 41,6 | 0,2 | 1 | 40,2 | 0,0 | 1 | 47,6 | 0,2 |
| TML | 10 | 47,7 | 0,0 | 10 | 41,9 | 0,2 | 10 | 40,2 | 0,0 | 10 | 47,1 | 0,4 |
| 5mC | 0,1 | 47,7 | 0,0 | 0,1 | 41,9 | 0,2 | **0,1** | **40,2** | **0,0** | 0,1 | 47,9 | 0,2 |
| 5mC | 1 | 47,7 | 0,0 | 1 | 41,9 | 0,2 | **1** | **40,5** | **0,0** | 1 | 47,9 | 0,2 |
| 5mC | 10 | 47,7 | 0,0 | 10 | 41,6 | 0,2 | **10** | **40,8** | **0,0** | 10 | 47,4 | 0,0 |
| C | - | 45,5 | 0,2 | - | 42,2 | 0,2 | - | 40,3 | 0,2 | - | 44,5 | 0,3 |
| m6A | 0,1 | 45,3 | 0,0 | 0,1 | 42,3 | 0,0 | 0,1 | 40,2 | 0,0 | 0,1 | 44,6 | 0,6 |
| m6A | 1 | 45,3 | 0,0 | 1 | 41,7 | 0,0 | 1 | 39,6 | 0,0 | 1 | 44,3 | 0,2 |
| m6A | 10 | 45,3 | 0,4 | 10 | 42,0 | 0,0 | 10 | 39,9 | 0,0 | 10 | 44,3 | 0,2 |
| 7mG | 0,1 | 45,3 | 0,0 | **0,1** | **42,3** | **0,0** | 0,1 | 40,2 | 0,0 | 0,1 | 44,1 | 0,4 |
| 7mG | 1 | 45,3 | 0,0 | **1** | **42,3** | **0,0** | 1 | 40,2 | 0,0 | 1 | 44,4 | 0,0 |
| 7mG | 10 | 45,6 | 0,0 | **10** | **41,7** | **0,0** | 10 | 40,2 | 0,0 | 10 | 44,6 | 0,2 |

C, control with 2-8 μM protein without ligand; sDMA, symmetrically methylated arginine; aDMA, asymmetrically methylated arginine, MML, mono-methylated lysine; DML, di-methylated lysine (DML); TML, tri-methylated lysine; 5mC, 5-methyl-cytosine; 6mA, 6-methyl-adenosine; 7mG, 7-methyl-guanosine triphosphate. Each ligand was tested at three concentrations; the standard deviation (S.D.) was calculated from duplicates. $\Delta T_m$s indicative of binding are highlighted in bold. The temperature differences were considered significant (shown in bold) if the direction of the temperature change was consistent over the three ligand concentrations and the difference between the melting temperatures at 10 mM and 0 mM ligand, with their error range, was higher than 0.5 °C.

**Table S2. Aub, Vasa, Piwi and Qin derived peptides carrying aDMA or sDMA marks tested for binding to Q-eTud1–5.**

| sDMA-peptide* | sequence |
|---|---|
| Aub11** | PVIARGRGR*** |
| Aub13 | IARGRGRGK |
| Aub15 | RGRGRGRKP |
| *Ago4* | MSGRGNLL |
| *Ago68* | ISVGRGRAR |
| *Ago70* | VGRGRARLI |
| Piwi7 | MADDQGRGRRRPL |
| Qin1148 | EIHVTGRGRTENS |
| Vasa17 | VDTRGARGGDWSD |
| Vasa65 | GRIGGGRGGGAGG |
| Vasa588 | RVGNNGRATSFFD |
| Vasa644 | GVDVRGRGNYVGD |

| aDMA-peptide**** | sequence |
|---|---|
| Vasa17 | VDTRGARGGDWSD |
| Vasa65 | GRIGGGRGGGAGG |
| Vasa73 | GGAGGYRGGNRDG |
| Vasa94 | EGERDFRGGEGGF |
| Vasa101 | GGEGGFRGGQGGS |
| Vasa108 | GGQGGSRGGQGGS |
| Vasa115 | GGQGGSRGGQGGF |
| Vasa122 | GGQGGFRGGEGGF |
| Vasa150 | RLDREERGGERRG |
| Vasa163 | RLDREERGGERGE |
| Aub382 | PTDKNIRGGNDQA |

*The interaction of the peptides carrying the sDMA mark with Q-eTud1–5 was probed by TSA and with Q-eTud3 and Q-eTud5 by NMR. The binding of the peptides derived from Ago3 (in cursive) was tested only in combination with Q-eTud3 and Q-eTud5 by NMR. In the NMR experiments the protein concentration was 50-200 μM and the peptide was added in 5- or 10-fold excess.
**The peptides are named by the name of the protein to which they belong followed by the number of the amino acid carrying the DMA mark in the full-length protein.
*** The underlined arginine carries the DMA modification.
**** The interaction of the peptides carrying the sDMA mark with Q-eTud1–5 was probed by TSA.

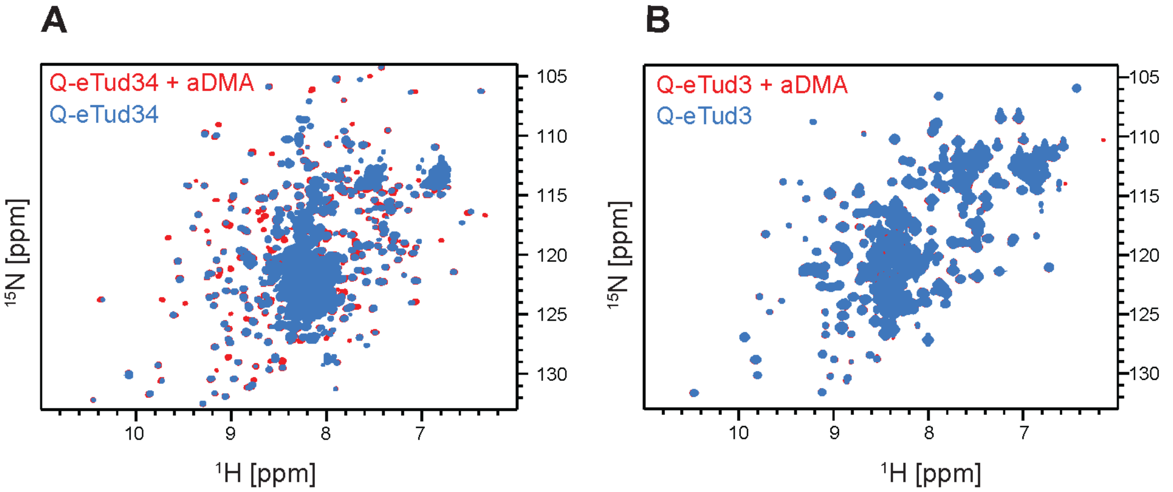


**Figure S1. Related to Figure 1: $^{1}$H,$^{15}$N HSQC spectra of Q-eTud34 and Q-eTud3. A.** Overlay of $^{1}$H-$^{15}$N HSQC spectra of 50 µM Q-eTud34 in isolation (blue) and in the presence of 2.5 mM aDMA (red). The appearance of CSPs confirms that Q-eTud34 binds aDMA. The spectra were recorded at 293 K and 850 MHz $^{1}$H frequency. **B.** Overlay of $^{1}$H-$^{15}$N HSQC spectra of 400 µM Q-eTud3 in isolation (blue) and in the presence of 6 mM aDMA (red). The absence of CSPs shows that Q-eTud3 does not bind aDMA. The spectra were recorded at 293 K and 600 MHz$^{1}$H frequency.

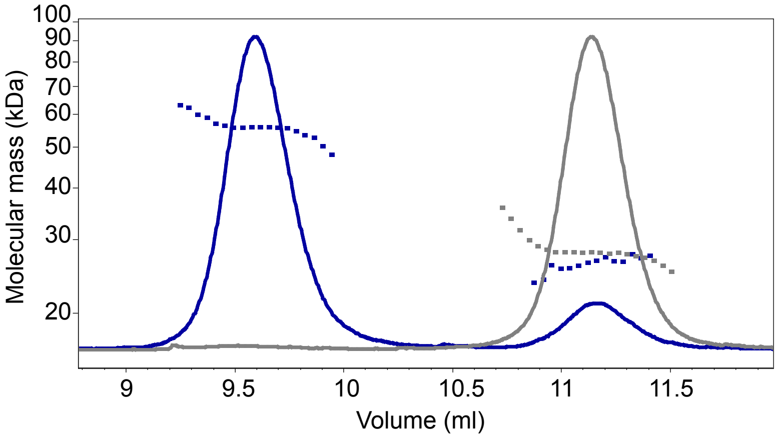


**Figure S2. Related to Figure 3: Q-eTud3 dimerizes.** Multi-angle light scattering profile of the purified size-exclusion chromatography peak corresponding to monomeric Q-eTud3 (gray) and of the peak corresponding to dimeric Q-eTud3 (blue). The elution volume on the x-axis refers to the size-exclusion chromatography run coupled with the multi-angle light scattering measurement. The blue trace shows that Q-eTud3d partially dissociates into monomers, as indicated by the appearance of a second peak at 11.2 ml, next to the main peak at 9.6 ml.

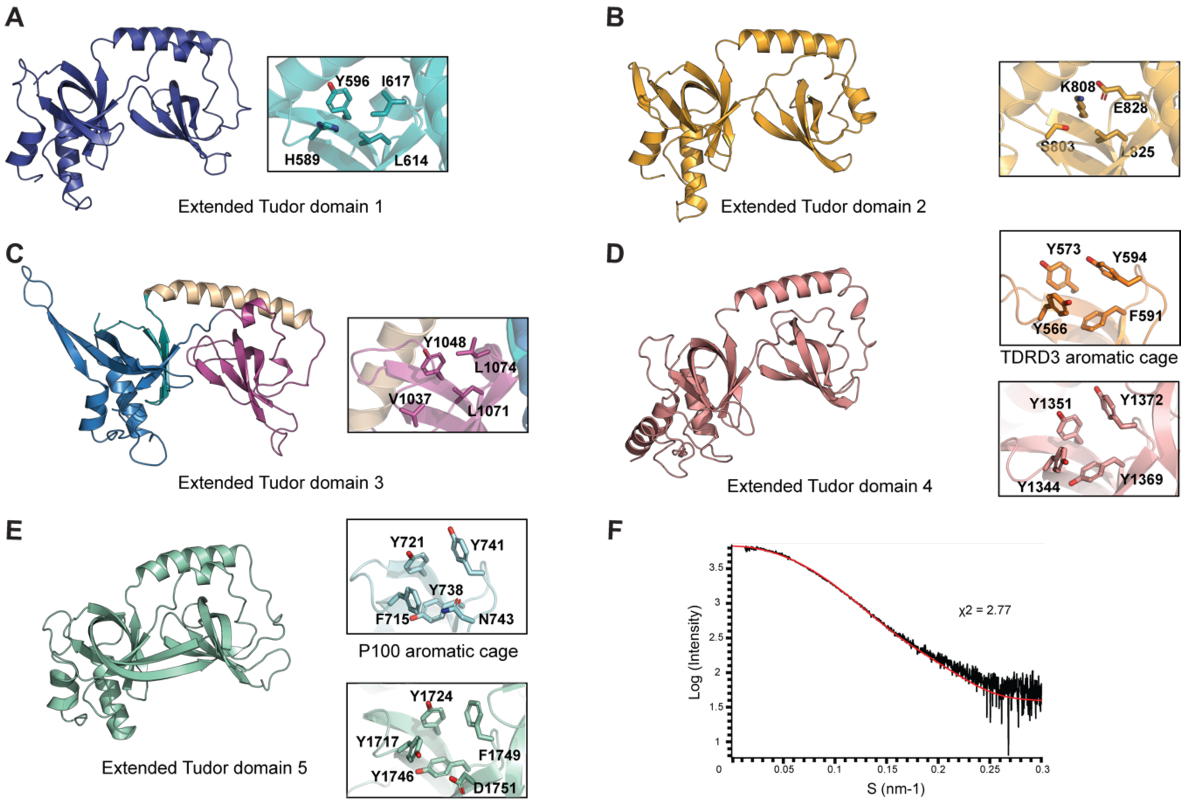


**Figure S3. Related to Figure 2: Homology models and structure of Qin's five extended Tudor domains. A.** Homology model of Q-eTud1. **B.** Homology model of Q-eTud2. **C.** X-ray structure of Q-eTud3. SN domain, blue; Tudor domain, magenta; connecting helix, yellow. **D.** Homology model of Q-eTud4. **E.** Homology model of Q-eTud5. The panels on the right show an expansion of the putative aromatic cage of all Qin Tudor domains. The aromatic cage is absent in Q-eTud1–3. The aromatic cage of Q-eTud4 (D, middle panel) resembles that of TDRD3 (PDB code 2lto) (D, right panel) (Sikorsky et al., 2012), which is known to bind aDMA. The aromatic cage of Q-eTud5 (E, middle panel) resembles that of P100 (PDB code 2o4x) (E, right panel) (Shaw et al., 2007), which is known to bind sDMA. **F.** Comparison of the experimental SAXS curve (black) of Q-eTud5, acquired with 8 mg/ml protein dissolved in buffer (25 mM potassium phosphate, 250 mM potassium chloride, 5 mM DTT, pH 6.0) at 293 K, with the theoretical curve (red) predicted from the homology model of panel E using Crysol (Svergun et al., 1995).

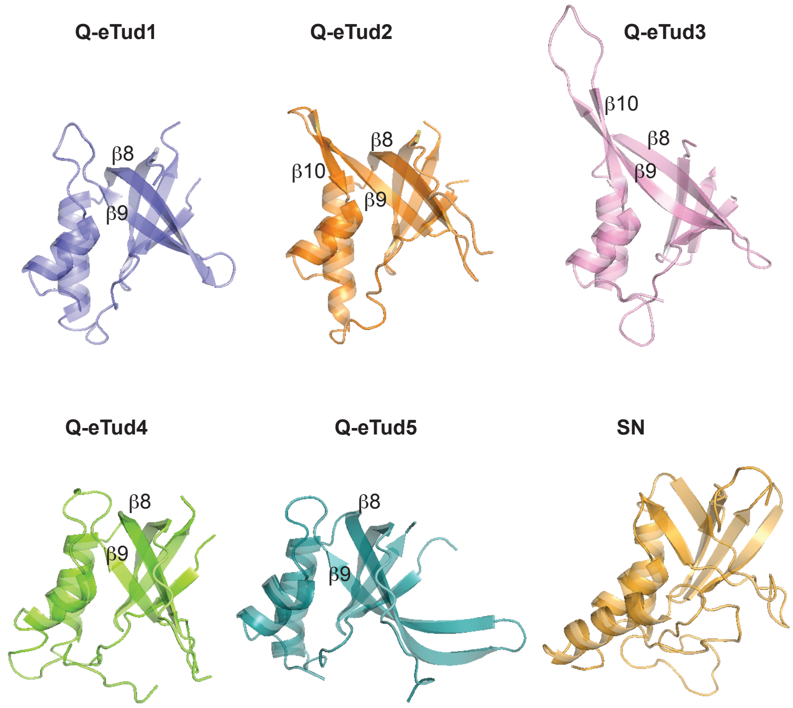


**Figure S4. Related to Figure 2: Structural differences of the SN domains of Q-eTud1–5.** The central β-sheet of the SN domains of QeTud1–5, consisting of β8 and β9, is longer than that of the SN domain of Staphylococcal Nuclease. β9 is extended either at the C-terminus (Q-eTud2 and Q-eTud3) or at the N-terminus (Q-eTud5), which yields to a prominent β-strand–loop–β-strand protuberance in Q-eTud3 and Q-eTud5.

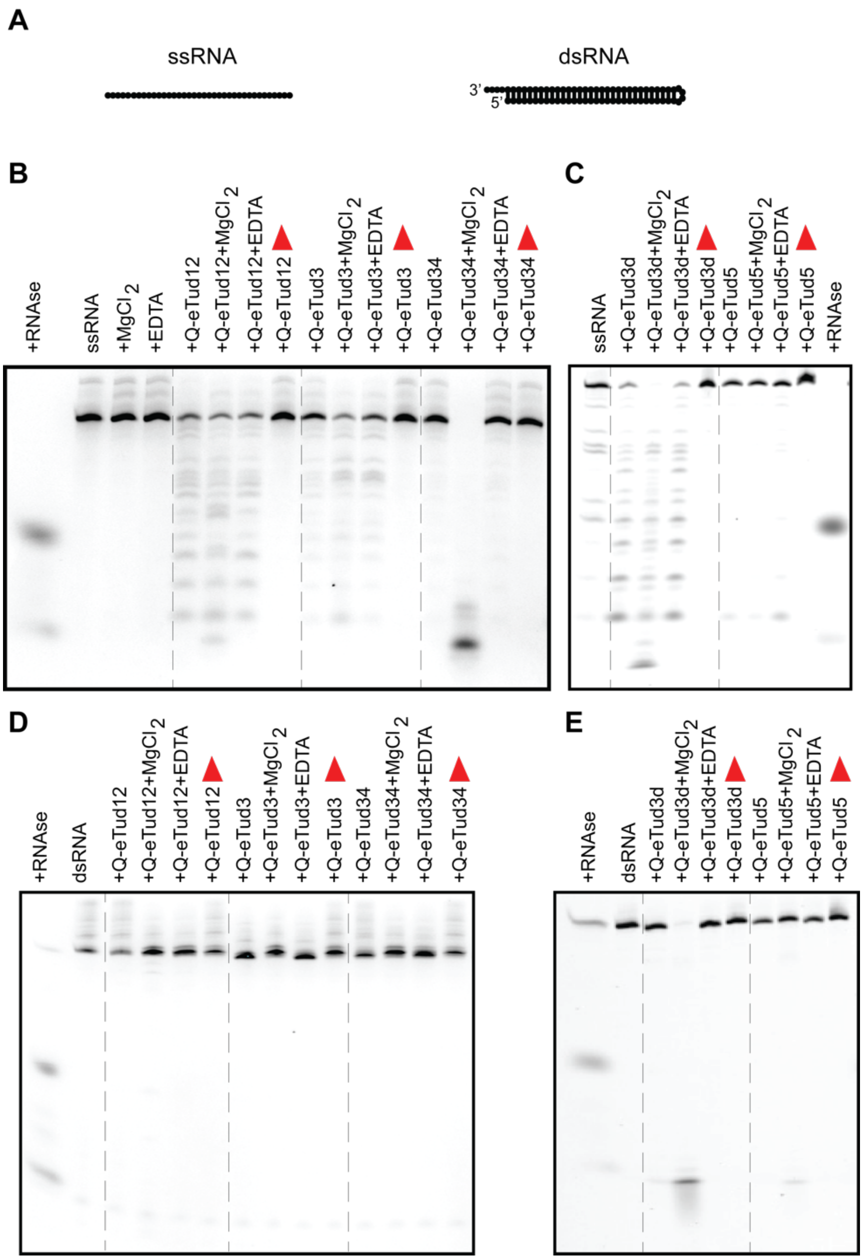


**Figure S5**. **Related to Figure 3: Q-eTud proteins are not active on dsRNA.** Q-eTud12, Q-eTud3, Q-eTud34 and Q-eTud5 were assessed for RNase activity against single-stranded (ss) and double-stranded (ds) RNA 65 and 66 nucleotides long, respectively. The assays were carried in buffer containing 25 mM HEPES potassium salt pH 7.0, 125 mM KCl, 1 mM DTT, 0.5 μM RNA and 40 μM protein. Each Q-eTud construct was assayed in the presence and in the absence of either $Mg^{2+}$ or EDTA (both at 10 mM). **A.** Secondary structures of the ssRNA and dsRNA. **B–E.** Denaturing PAGE showing ~0.5 μM of the 3' end fluorescent labeled RNAs after 5.5 hours of incubation at 30 °C with the Q-eTud construct indicated above each lane. The control lanes, containing RNA incubated with heat-denatured protein buffer (25 mM HEPES potassium salt pH 7.0, 125 mM KCl, 1 mM DTT), are labeled with a red triangle. As a comparison of nuclease activity, 0.15 mU of RNaseA were added to the same RNA in the first lane. Q-eTud3m and Q-eTud3d indicate the monomeric and dimeric forms of the protein, respectively. The dashed lines separate the lanes containing different Q-eTud constructs.

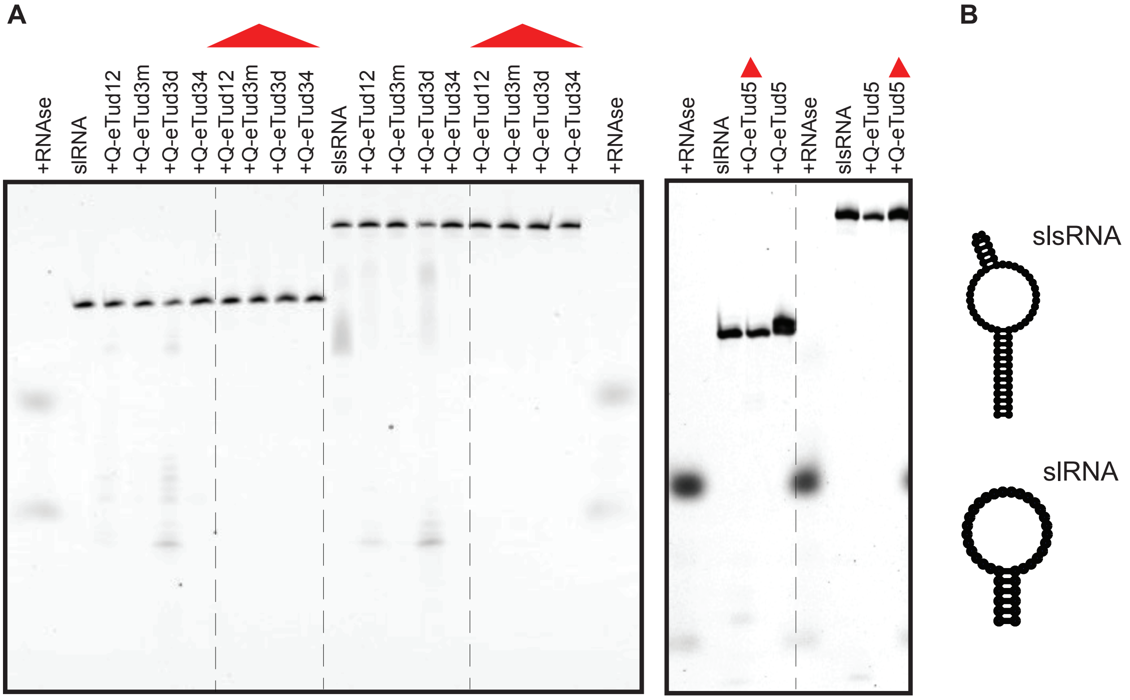


**Figure S6. Related to Figure 3: Q-eTuds are not active on stem-loop RNAs. A.** Q-eTud12, Q-eTud3m (monomeric form), Q-eTud3d (dimeric form), Q-eTud34 and Q-eTud5 were assessed for RNase activity against stem loop (sl) and stem-loop-stem (sls) RNAs 39 and 67 nucleotides long, respectively. The assays were carried in buffer containing 25 mM HEPES potassium salt pH 7.0, 125 mM KCl, 1 mM DTT, 10 mM $MgCl_2$, 0.5 µM RNA and 40 µM protein. The denaturing PAGE shows ~0.5 µM of the 3' end fluorescent labeled RNAs after 5.5 hours of incubation at 30 °C with the Q-eTud construct indicated above each lane. The control lanes, containing RNA incubated with heat-denatured protein in the same buffer, are labeled with a red triangle. The RNA construct used in the assay is indicated above the control lane. As a comparison of nuclease activity, 0.15 mU of RNaseA were added to the same RNA in the first lane. The dashed lines separate the lanes containing either different RNAs or the heat-denatured proteins. **B.** Predicted secondary structures of the slRNA and slsRNA.

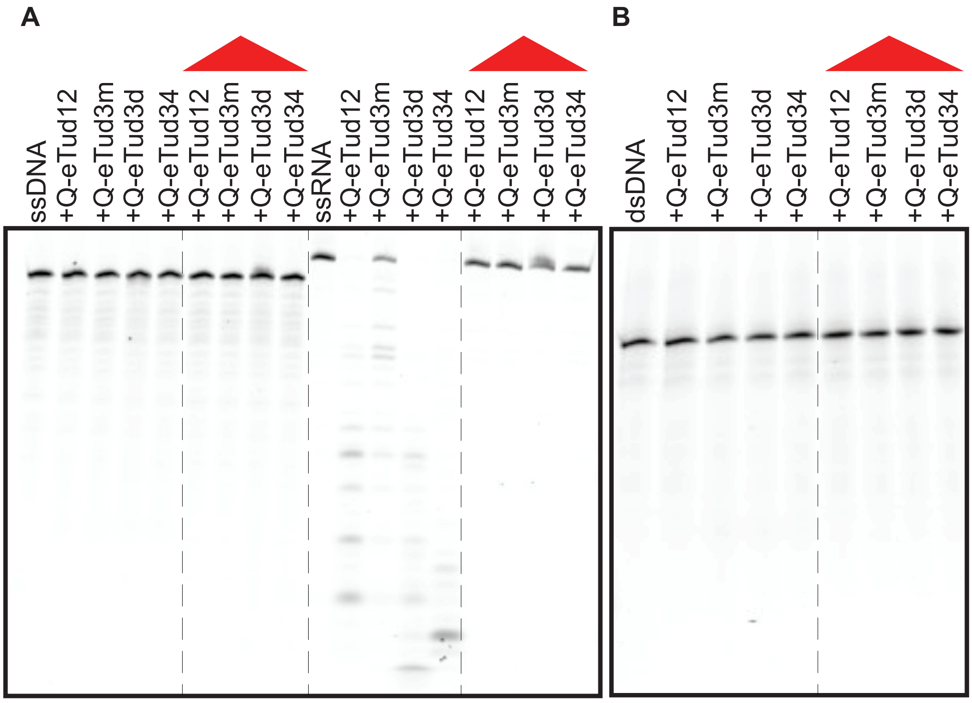


**Figure S7. Related to Figure 3: Qin does not have DNase activity. A and B.** The DNase and RNase assays were conducted in buffer containing 25 mM HEPES potassium salt pH 7.0, 125 mM KCl, 1 mM DTT, 10 mM $MgCl_2$, 40 μM protein and 0.5 μM of either ssDNA or ssRNA (A) or dsDNA (B). The denaturing PAGE shows ~0.5 μM of the 3' end fluorescent labeled DNA or RNA after 5.5 hours of incubation at 30 °C with the Q-eTud construct indicated above each lane. The control lanes, containing RNA or DNA incubated with heat-denatured protein in the same buffer, are labeled with a red triangle The DNA or RNA construct is indicated above the control lane. The dashed lines separate the lanes containing either different nucleic acids or the heat-denatured proteins.

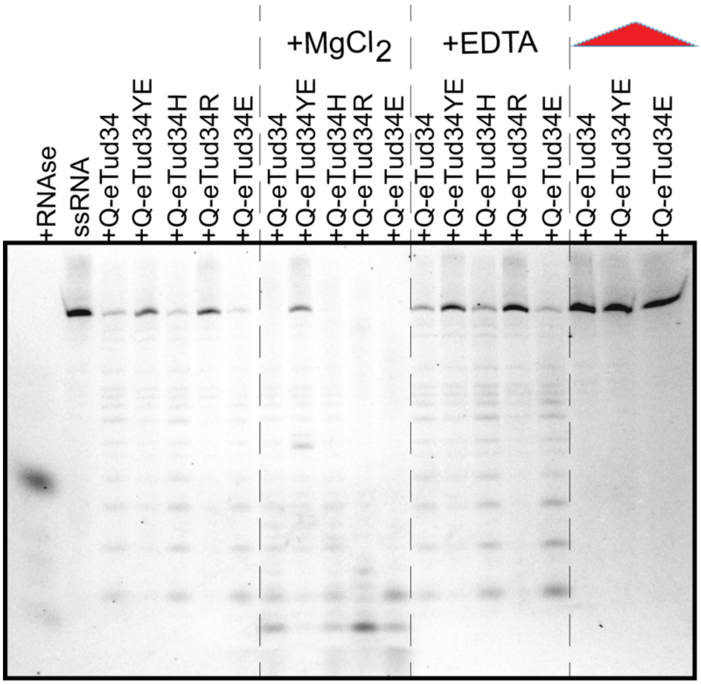


**Figure S8. Related to Figure 5: RNase assay of Q-eTud4 mutants in the Q-eTud34 construct in the presence or absence of magnesium.** The mutants tested were Q-eTud34 Y1292A/E1293A (Q-eTud34YE), H1402A (Q-eTud34H), R1405A (Q-eTud34R) and E1415A (Q-eTud34E). The RNase activity of the Q-eTuds was evaluated upon incubation for 5.5 hours at 30 °C of ~0.5 µM 3' fluorescently-labeled ssRNA with 40 µM Q-eTud34 either in buffer (buffer composition: 25 mM HEPES potassium salt pH 7.0, 125 mM KCl, 1 mM DTT) or in buffer in the presence of either 10 mM $MgCl_2$ or 10 mM EDTA. The control lanes, containing RNA incubated with heat-denatured protein in buffer, are labeled with a red triangle. The dashed lines separate either lanes in different buffer conditions or lanes containing the heat-denatured proteins.
